# Sequential predictive e-diagnostics for hidden Markov models of animal movement

**DOI:** 10.64898/2026.07.07.737005

**Authors:** Aurélien Nicosia

## Abstract

Hidden Markov models are standard for inferring behavioural states from animal movement data, but checking whether a fitted latent-state model predicts held-out movement well remains difficult. We develop sequential predictive e-diagnostics that evaluate a fitted movement HMM as a generator of validation trajectories. Each diagnostic specifies a predictable alternative density, and its ratio to the fitted model’s observable one-step predictive density defines an e-value increment. The denominator is obtained by filtering over latent states, not by conditioning on a decoded path. Under a fixed train/validation protocol, the cumulative product is an e-process, giving anytime-valid thresholds under optional stopping and predictable switching. The construction extends to weighted and state-localized evidence, feature-level circular-linear checks, and blockwise summaries. Controlled simulations show calibration under the fitted-generator null and sensitivity to targeted misspecifications. A leave-one-animal-out elk case study illustrates pooled, individual-specific and state-localized predictive model criticism in a standard movement-HMM workflow.

## 1 Introduction

Hidden Markov models (HMMs) are central in animal movement analysis when movement is viewed as the observable output of unobserved behavioural states. They sit within the broader state-space tradition for linking telemetry to latent behavioural and spatial dynamics (Patterson et al., 2017). In common step-angle models, latent states generate step lengths and turning angles and are interpreted as resting, foraging, exploratory movement or directed travel. This mixture random-walk construction (Morales et al., 2004) is now standard in movement ecology and ecological state-dynamics applications (Langrock et al., 2012; McClintock et al., 2020; Zucchini et al., 2016), with implementations in `moveHMM` (Michelot et al., 2016) and `momentuHMM` (McClintock and Michelot, 2018).

As a motivating example, consider the elk telemetry data distributed with `moveHMM` (Michelot et al., 2016; Morales et al., 2004), in which four animals are summarized by step lengths and turning angles. A practitioner fitting a multi-state HMM must decide how many behavioural states to retain and whether the fitted model predicts animals not used for fitting. AIC and BIC rank candidate models, but they do not say whether the selected model predicts a held-out trajectory well, nor where it fails. Section 7 returns to these data under leave-one-animal-out (LOAO) validation.

Existing assessment tools address parts of this problem. Penalized likelihood criteria, cross-validated likelihood and pragmatic workflows guide state-number selection (Langrock et al., 2017). Probability integral transform and pseudo-residual diagnostics use one-step forecast distributions (Dunn and Smyth, 1996; Zucchini et al., 2016), while HMM-specific residual and goodness-of-fit checks provide broader misspecification tools (Buckby et al., 2020). The diagnostics proposed here complement rather than replace these methods.

What remains less developed is validation that is predictive, sequential and explicit about latent-state uncertainty. Diagnostics may be chosen after seeing the data, decoded states may be treated as observed despite being model-dependent reconstructions, and a high fitted likelihood or plausible state interpretation need not imply good prediction of held-out movement, dwell times, directional persistence or spatial summaries.

This paper proposes a predictive alternative. The fitted HMM is evaluated as a sequential generator of a validation trajectory. At each validation time, the null model supplies the observable one-step predictive density, a diagnostic alternative supplies a predictable competing density, and their ratio is a predictive e-value increment. The cumulative product is an e-process under the fixed null generator and can be monitored with Ville-type thresholds (Ville, 1939; Howard et al., 2020). The contribution is not simply to import e-values into movement ecology, but to build valid diagnostics around the observable predictive law of a latent-state movement model. The construction connects prequential evaluation (Dawid, 1984) with e-values, safe testing and anytime-valid inference (Vovk and Wang, 2021; Ramdas et al., 2023; Grünwald et al., 2024). Formal notation begins in Sections 2 and 3.

The paper makes four methodological contributions. It constructs predictive e-processes for fixed movement HMMs evaluated on held-out validation sequences; clarifies why validity requires the observable predictive density obtained by filtering over latent states, not conditioning on a Viterbi or otherwise decoded path; develops movement-oriented extensions for predictable mixtures and switching, localization, feature-level diagnostics and blockwise diagnostics; and evaluates the workflow through controlled simulations and an LOAO elk analysis. The guarantees are exact under a fixed train/validation protocol in which the null model and diagnostic alternatives are fixed before validation. Same-sample workflows remain exploratory unless additional sample splitting, cross-fitting or pre-specified conditioning arguments are used.

The paper next defines the validation protocol and observable predictive density (Section 2), gives the e-process construction (Section 3), develops movement-specific diagnostics (Section 4), treats split validation and composite nulls (Section 5), and presents simulations, the elk application and discussion.

## 2 Validation protocol and observable prediction for movement HMMs

### 2.1 Training information and validation filtration

Let (Ω, *A*) be the underlying measurable space. Let ℋ ⊆ *A* be the pre-validation sigma-field generated by the training data, fitted parameters, preprocessing choices, random seeds, and diagnostic-menu choices. All modelling choices in ℋ are fixed before validation.

The validation trajectory is indexed by *t* = 1, …, *T*. The observed movement variable is a measurable mapping *Y*_*t*_ : Ω → *Y*_*t*_, where *Y*_*t*_ is the observation space at time *t*. In the step-turning-angle setting,

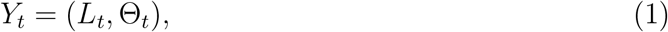

where *L*_*t*_ is a step length and Θ_*t*_ is a turning angle. Lowercase *y*_*t*_ = (*l*_*t*_, *θ*_*t*_) denotes a generic realized value of *Y*_*t*_. More generally, *Y*_*t*_ may include several movement streams or covariates observed at the same temporal resolution. The latent behavioural state is

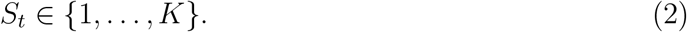

Here *K* is the number of states in the fitted HMM. The observable validation filtration is defined as

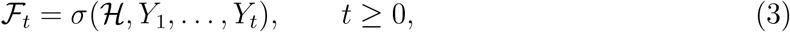

where ℱ_0_ = ℋ. We work with the raw discrete-time filtration, or with a universal completion when required under a composite family. A diagnostic rule is *predictable* if the object used at time *t* is ℱ_*t−*1_-measurable. Thus the diagnostic predictive alternative for *Y*_*t*_ must be chosen before *Y*_*t*_ is revealed.

Throughout, *M*_0_ denotes the fitted null generator after training, and ℙ_0_ denotes the conditional validation law induced by *M*_0_ after conditioning on ℋ. Under this law, the validation sequence has one-step predictive density *p*_0_(*y*_*t*_ | ℱ_*t−*1_). All e-process guarantees are conditional on ℋ, and unconditional guarantees follow by iterated expectation. This separates fitted-generator validation from parameter-uncertainty questions.

The notation used throughout the theoretical setup is collected in a summary table in the supplementary material.

### 2.2 Movement HMM and predictable covariates

Conditional on ℋ, the null model *M*_0_ is a fixed *K*-state HMM with initial distribution ***π***_0_, transition matrix **Γ**_0_ and state-dependent observation densities. Let *X*_*t*_ be the vector of covariates available at step *t*. Write

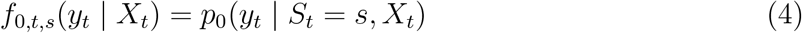

for the state-dependent density in state *s*. We assume that covariates *X*_*t*_ used to predict *Y*_*t*_ are ℱ_*t−*1_-measurable. Environmental covariates evaluated at the animal’s starting location at time *t* − 1 are therefore predictable, whereas covariates evaluated at the arrival location of step *t* generally depend on *Y*_*t*_ and are not predictable unless evaluated under the predicted step distribution. Below, 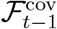 denotes the predictable covariate information used in the observation density. In a common movement specification,

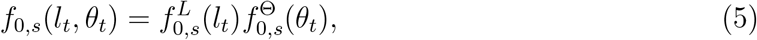

for example with Gamma step lengths and von Mises turning angles. The factorization in (5) is a modelling assumption, not a requirement of the e-process construction.

The HMM formulation also allows covariate-dependent transition probabilities, time-varying state-dependent parameters and multivariate observations, provided the one-step predictive density is well-defined and any covariates used to predict transitions Γ_0,*t*_(*r, s*) or observation densities are predictable.

Three validation targets should be separated: the fixed fitted generator, the true parametric HMM with estimated parameters, and a composite null family. This paper focuses primarily on the first target, asking whether the specific fitted HMM resulting from training generates the validation trajectory. Parameter uncertainty and composite-null validity require additional treatment; a summary table of these targets is given in the supplementary material.

### 2.3 Observable predictive density by filtering

The central object is the one-step-ahead observable predictive density. Since *S*_*t*_ is latent, this density must marginalize over the predictive distribution of the state:

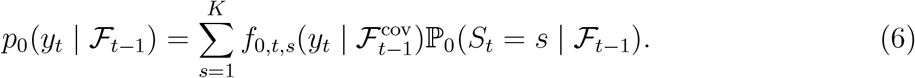

This is the density of the next movement observation under the fitted HMM, conditional only on the observed past and the training information.

Let

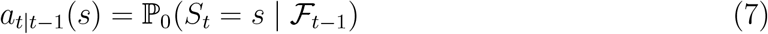

denote the predictive state probabilities before observing *Y*_*t*_. The recursion is initialized with the stationary or fitted initial distribution, *a*_1|0_(*s*) = *π*_0_(*s*). With multiple validation individuals, the filter is restarted for each held-out individual *i* using *a*_*i*,1|0_(*s*) = *π*_0,*i*_(*s*), where *π*_0,*i*_ is the fitted initial distribution for that individual or fold. Then

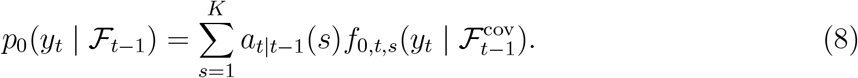

After observing *Y*_*t*_ = *y*_*t*_, the filtered probabilities are

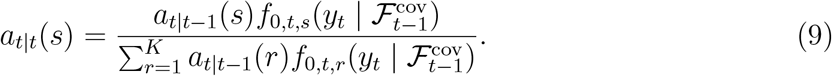

The next predictive state probabilities are

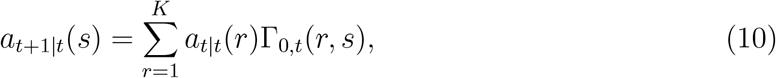

where Γ_0,*t*_(*r, s*) = ℙ_0_(*S*_*t*+1_ = *s* | *S*_*t*_ = *r*, ℱ_*t*_) defines the transition convention. If transitions depend on covariates, Γ_0,*t*_ must be ℱ_*t*_-measurable. Spatial covariates computed from the animal’s arrival location are predictable only if determined before observing *Y*_*t*_, or if the diagnostic targets the joint distribution of movement and spatial context. In the time-homogeneous case, Γ_0,*t*_ ≡ Γ_0_; with covariate-dependent transitions, the filter should be initialized using *π*_0_ or another pre-specified distribution.

## 3 Sequential predictive e-diagnostics

### 3.1 Diagnostic alternatives as predictable densities

A diagnostic alternative is represented by a predictable Markov kernel or sub-Markov kernel *Q*_*t*_. When represented by a density or subdensity *q*_*t*_(*y* | ℱ_*t−*1_) with respect to a common dominating measure, *y* is a generic element of *Y*_*t*_, and the map (*ω, y*) → *q*_*t*_(*y* | ℱ_*t−*1_)(*ω*) must be measurable with respect to past information and the observation sigma-field on *Y*_*t*_. The subscript *t* allows the diagnostic to depend on the observed past. The key requirement is predictability: the rule defining *q*_*t*_ must be fixed before observing *Y*_*t*_.

Examples include:

- a *K* + 1 state HMM used to diagnose whether the null HMM has too few states;
- a more flexible angular distribution used to diagnose misspecified turning angles;
- a joint step-angle model used to diagnose residual dependence between *L*_*t*_ and Θ_*t*_ within behavioural states;
- a duration-aware alternative, such as a hidden semi-Markov model (HSMM), used to diagnose non-geometric state dwell times;
- a flexible predictive model trained on independent data and used as a diagnostic competitor to the fitted HMM.

These alternatives are diagnostic bets. They need not be interpreted as true data-generating mechanisms.

Throughout, all predictive distributions and diagnostic bets are represented by densities with respect to a common dominating measure *µ*_*t*_ on the observation space at time *t*. This can be Lebesgue measure for continuous step-angle observations, counting measure for discrete observations or a product measure for mixed data. The convention is that ratios are evaluated only on the support of the null predictive density; outside that support the null event has probability zero, and we adopt the support convention that *q*_*t*_(*y* | ℱ_*t−*1_) = 0 whenever *p*_0_(*y* | ℱ_*t−*1_) = 0. Diagnostic bets *q*_*t*_ may be fixed before validation, or updated sequentially online using past validation observations, provided the update rule is predictable.

### 3.2 Predictive e-value increments and e-processes

#### **Definition 1** (Predictive e-value increment and process).

For *t* = 1, …, *T*, define

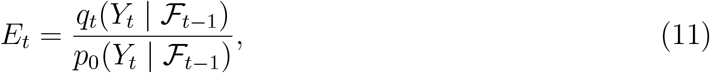

whenever the ratio is well-defined under *M*_0_. The associated cumulative process is

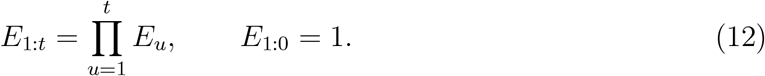

Equivalently,

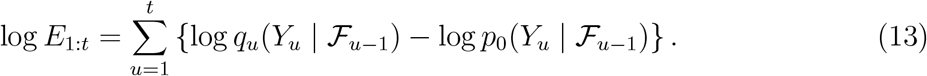

#### **Theorem 1** (Predictive e-process for a fixed HMM).

*Assume that, conditional on* ℋ, *the validation sequence is generated under the null HMM M*_0_, *and that p*_0_(*y* | ℱ_*t−*1_) *is the corresponding observable predictive density with respect to µ*_*t*_. *Let q*_*t*_(*y* | ℱ_*t−*1_) *be an* ℱ_*t−*1_*-measurable diagnostic subdensity satisfying*

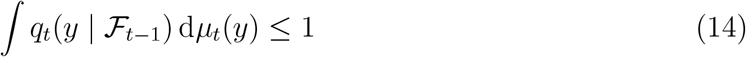

*almost surely. We assume either q*_*t*_ ≪ *p*_0_ *almost surely, or use the convention E*_*t*_ = 0 *on* {*y* : *p*_0_(*y* | ℱ_*t−*1_) = 0} *under the validation law* ℙ_0_. *Then*

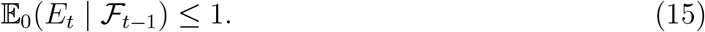

*Consequently*, (*E*_1:*t*_)_0*≤t≤T*_ *is a nonnegative supermartingale under M*_0_. *If q*_*t*_(· | ℱ_*t−*1_) *is a density assigning all its mass to the support of p*_0_(· | ℱ_*t−*1_), *then equality holds and* (*E*_1:*t*_)_0*≤t≤T*_ *is a martingale*.

#### Proof

Conditional on ℱ_*t−*1_, *Y*_*t*_ has density *p*_0_(*y* | ℱ_*t−*1_) under *M*_0_. Hence, with the stated support convention,

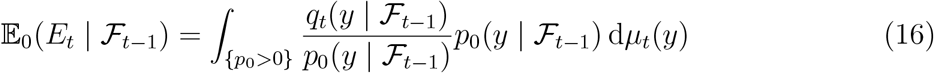

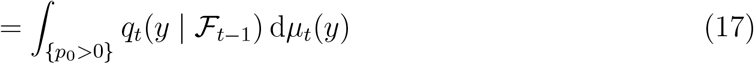

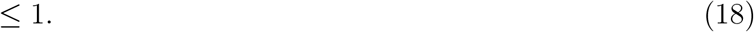

Therefore *E*_0_(*E*_1:*t*_ | F_*t−*1_) = *E*_1:*t−*1_*E*_0_(*E*_*t*_ | ℱ_*t−*1_) ≤ *E*_1:*t−*1_, so the cumulative process is a nonnegative supermartingale. Equality holds when *q*_*t*_ is a density supported by the null support.

#### **Corollary 2** (Optional stopping and anytime-valid monitoring).

*Under the conditions of*

#### *Theorem 1, for any α* ∈ (0, 1),

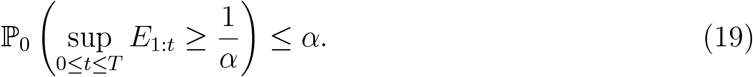

*Equivalently*,

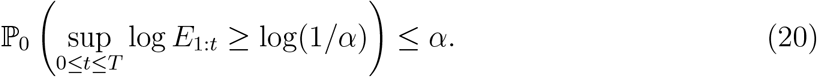

*Moreover, for any stopping time τ with respect to* (F_*t*_), *the stopped process E*_1:*τ∧T*_ *is an e-value under M*_0_, *meaning E*_0_(*E*_1:*τ∧T*_) ≤ 1.

#### Proof

The maximal inequality is Ville’s inequality applied to the nonnegative supermartingale (*E*_1:*t*_)_0*≤t≤T*_; the logarithmic statement follows by monotonicity of the logarithm. For any stopping time *τ*, optional stopping at the bounded time *τ* ∧*T* gives *E*_0_(*E*_1:*τ∧T*_) ≤ *E*_1:0_ = 1.

The equivalent anytime p-value formulation is stated in the supplementary material.

### 3.3 Why filtering rather than decoding is essential

The proof of Theorem 1 requires the denominator to be the conditional density of the next observable variable given the observable past. In an HMM, this is the filtered marginal predictive density, not a density conditional on a decoded state.

#### **Proposition 3** (Validity is based on marginal observable prediction).

*For an HMM, the denominator in* (11) *is the marginal observable predictive density*

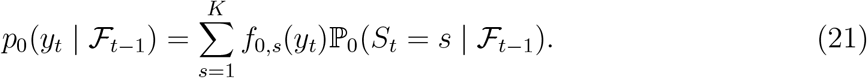

*The e-process validity does not require observing S*_*t*_ *and does not generally follow from replacing S*_*t*_ *by a decoded path Ŝ*_*t*_.

#### Proof

The conditional law of *Y*_*t*_ given ℱ_*t−*1_ is obtained by summing over the unobserved state *S*_*t*_ using predictive state probabilities. A decoded path *Ŝ* _*t*_ is a statistic produced by the fitted model and observed data; it does not enter this conditional law unless it is known to equal the true latent state, which is not the usual HMM setting.

The same issue arises for any data-dependent decoded state 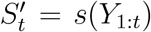: the denominator 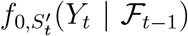 depends on the current observation through 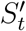 and need not give an expectation-bounded ratio. A two-state counterexample is given in Online Supplement S2.

#### Remark 1 (Decoded states are useful after, not before, validation).

A Viterbi path, local decoding or posterior state classification may help interpret a signal after validation. Such summaries are descriptive unless the localization weights are predictable; Section 4.2 gives a valid filtered-probability alternative.

## 4 Movement-specific diagnostic constructions

### 4.1 Predictable mixtures and switching

For a pre-specified menu of diagnostic alternatives, predictable mixtures remain valid. If *q*_*j,t*_(*y* | ℱ_*t−*1_), *j* = 1, …, *J*, are predictable diagnostic subdensities and *w*_*j,t*_ ≥ 0 are predictable weights with _*j*_ *w*_*j,t*_ ≤ 1, then

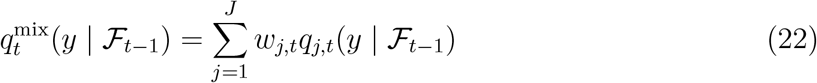

defines another valid diagnostic subdensity. Predictable switching is the special case with one nonzero weight.

This validity does not extend to products of diagnostics computed in parallel on the same observations. Unless a sequential conditional construction is justified, report individual e-processes and use weighted averages or weighted thresholds; see Online Supplement S3.

### 4.2 Predictable localization

Localization is obtained by predictable tempering. For any conditional e-value increment *E*_*t*_ and predictable weight *W*_*t*_ ∈ [0, 1],

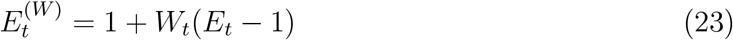

is again a valid conditional e-value increment. The weight may select seasons, habitats, individuals or management periods, provided it is chosen before the next response.

For behavioural-state localization, a valid soft weight is the filtered predictive probability

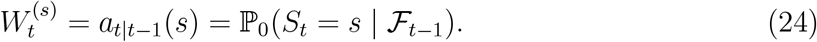

This localizes evidence to times where the fitted model predicts state *s* to be likely. Smoothed probabilities or decoded states may help post hoc interpretation, but are not predictable. The elk application uses this filtered-probability localization.

### 4.3 Feature-level and blockwise movement diagnostics

Many diagnostics target a lower-dimensional feature

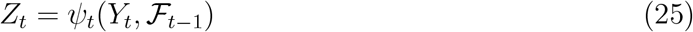

such as a circular residual, transformed step length, dependence residual or barrier-crossing indicator. If *g*_0,*t*_ is the null conditional law of *Z*_*t*_ and *g*_1,*t*_ is a predictable diagnostic subdensity, then

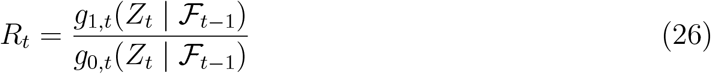

is again a conditional e-value increment. Rosenblatt transforms diagnose residual step-angle dependence, and blockwise summaries such as displacement, straightness, residence, barrier crossing or space use are valid when their null predictive law is known or independently estimated. Details are in Online Supplement S3.

### 4.4 Practical diagnostic menu for movement HMMs

Together these constructions cover common movement-HMM failures: missing behavioural states, angular misspecification, residual step-angle dependence, non-geometric dwell times, temporal nonstationarity, individual heterogeneity and blockwise spatial failures. A detailed catalogue is given in Online Supplement S7.

## 5 Split validation, cross-fitting, and composite nulls

The exact theory treats *M*_0_ as fixed at validation time, whereas applied movement models are fitted. This section separates conditional validity after training, cross-fitted e-values over independent validation units and conservative composite-null envelopes.

### 5.1 Conditional validity after training

#### **Theorem 4** (Split-sample conditional validity).

*Let* ℋ *contain the training data and all fitted parameters of the null and diagnostic alternatives. Suppose that, conditional on* ℋ, *the validation trajectory is generated from the fitted null predictive law p*_0_(· | ℱ_*t−*1_), *and suppose q*_*t*_ *is predictable with respect to* ℱ_*t−*1_ = H ∨ *σ*(*Y*_1_, …, *Y*_*t−*1_). *Then the process in Theorem 1 is an e-process conditional on* ℋ. *In particular*,

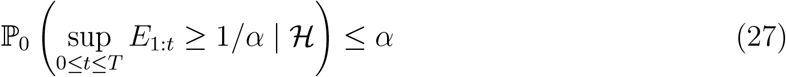

*almost surely, and the same bound holds unconditionally*.

#### Proof

Conditional on ℋ, the fitted densities are fixed and the validation filtration satisfies the assumptions of Theorem 1. The conditional bound follows from Corollary 2. Taking expectations over H gives the unconditional bound.

This theorem clarifies the interpretation of the first article’s target: the exact null is that the held-out validation sequence is generated by the fitted HMM. It is a diagnostic check of the fitted generator, not a claim that the parameter estimator has no uncertainty.

### 5.2 Independent validation units and cross-fitted averages

A natural movement design trains on some individuals and validates on held-out individuals. If each held-out individual yields an e-value, these e-values can be reported individually or combined by averaging.

#### **Proposition 5** (Cross-fitted average over validation units).

*Suppose E*^(*i*)^ *is an e-value under M*_0_ *for validation unit i, for i* = 1, …, *n, where each E*^(*i*)^ *may be constructed using a model fitted on data not including the validation responses of unit i. If w*_*i*_ ≥ 0 *and* _*i*_ *w*_*i*_ ≤ 1, *then*

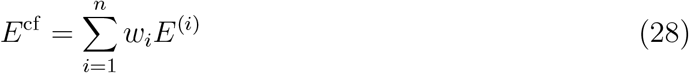

*is an e-value under M*_0_.

#### Proof

The proof is the same as for the parallel-diagnostics proposition in the supplementary material: *E*_0_*E*^cf^ = _*i*_ *w*_*i*_*E*_0_*E*^(*i*)^ ≤ 1. No independence among the e-values is needed for averaging. Independence or a suitable sequential conditional construction would be needed for products.

#### *Remark* 2 (Leave-one-individual-out products and joint nulls).

A leave-one-individual-out workflow may produce individually valid e-values, but the product across individuals is not automatically valid because the fitted models may use overlapping training sets and the e-values are computed in parallel. Specifically, conditional on the training set for fold *i, E*^(*i*)^ is valid for the hypothetical validation law generated by the fold-specific fitted model. The average *E*^cf^ is valid if all fold-specific validation laws are embedded in a common joint null under which each *E*^(*i*)^ has expectation at most one. Alternatively, leave-one-animal-out (LOAO) cross-validation should be presented as a practical cross-predictive diagnostic rather than an exact global test, unless a coherent joint null is specified. The weighted average in Proposition 5 is a safer default unless a sequential data-collection or sample-splitting design is used.

### 5.3 Conservative composite-null envelopes

The fitted-generator null is the most transparent target for model criticism. When a single process valid for every parameter value in a null family is required, a conservative envelope construction replaces the denominator by the pointwise supremum of the null predictive densities over the family. The resulting ratio is an e-process that is uniformly valid over the family, at the cost of conservatism, and is practical mainly for finite or compactly restricted families rather than general HMM parameter spaces. The formal statement, its proof, and a discussion of its cost are given in the supplementary material, together with the finite-family simulation S12.

### 5.4 What the exact guarantee does and does not claim

The exact guarantee is conditional on a fixed train/validation protocol. Once H fixes the fitted null generator, preprocessing, covariate choices and diagnostic menu, the validation sequence is used only through predictable density ratios. The result does not fully solve parametric estimation uncertainty for a true unknown HMM, and it does not make same-sample exploratory workflows confirmatory. Such workflows remain useful for model criticism, but confirmatory use requires sample splitting, cross-fitting, pre-specification or another valid conditioning argument. Composite-null envelopes provide uniform validity over a family, but they are conservative. Additional log-score and growth results are given in Online Supplement S2.

## 6 Simulation study

The simulations check calibration and targeted sensitivity in a reproducible R workflow. All scenarios use simulated step lengths and turning angles, forward filtering for the observable HMM predictive density, and cumulative log e-processes monitored against log(1*/α*); they are controlled diagnostic checks, not benchmarks of HMM-fitting algorithms. Several alternatives are fixed or oracle-style diagnostic competitors, so their role is to verify the behavior of the e-diagnostic construction under known misspecification rather than to compare estimation procedures.

Table 1 summarizes the seven scenarios retained in the main article: S1 and the null runs in S6 to S8 check calibration; S3 to S6 target an insufficient number of states, angular misspecification, residual step-angle dependence and non-geometric durations; and S7 to S8 check predictable localization, mixtures and switching.

**Table 1:**
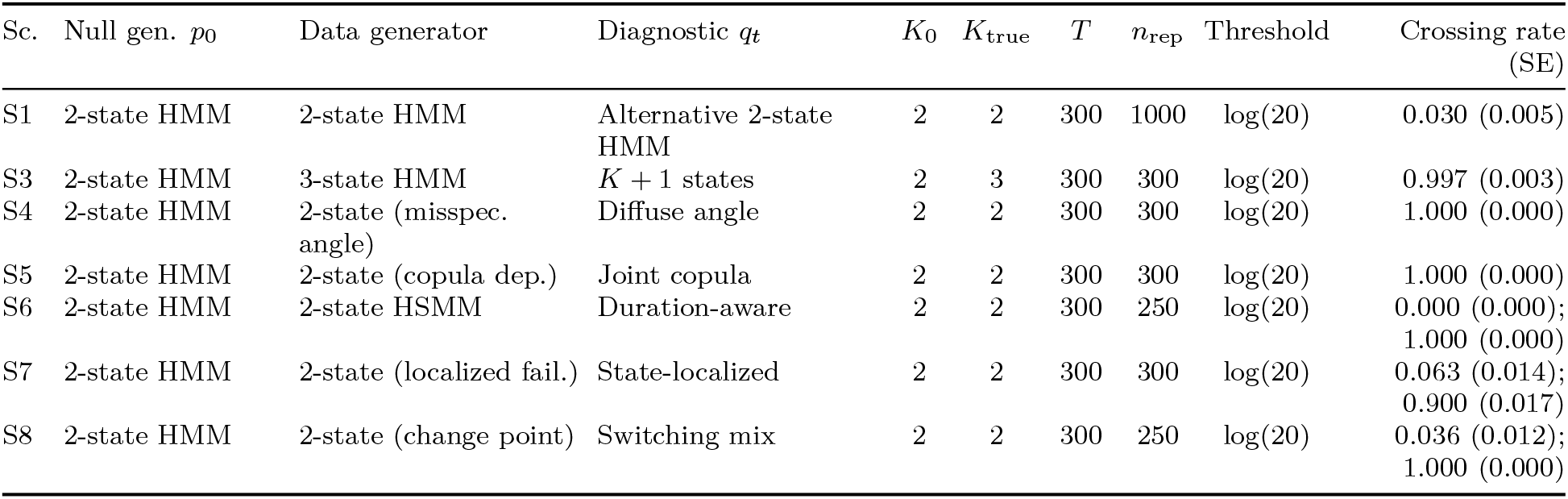
Simulation scenarios retained in the main article. When two crossing rates are shown, the first is the null calibration run and the second is the targeted alternative. For S6, these correspond to an HMM null calibration run and the HSMM alternative, both at the blockwise updated threshold log(20).

| Sc. | Null gen. $p_0$ | Data generator | Diagnostic $q_t$ | $K_0$ | $K_{\text{true}}$ | $T$ | $n_{\text{rep}}$ | Threshold | Crossing rate (SE) |
| --- | --- | --- | --- | --- | --- | --- | --- | --- | --- |
| S1 | 2-state HMM | 2-state HMM | Alternative 2-state HMM | 2 | 2 | 300 | 1000 | $\log(20)$ | 0.030 (0.005) |
| S3 | 2-state HMM | 3-state HMM | $K + 1$ states | 2 | 3 | 300 | 300 | $\log(20)$ | 0.997 (0.003) |
| S4 | 2-state HMM | 2-state (misspec. angle) | Diffuse angle | 2 | 2 | 300 | 300 | $\log(20)$ | 1.000 (0.000) |
| S5 | 2-state HMM | 2-state (copula dep.) | Joint copula | 2 | 2 | 300 | 300 | $\log(20)$ | 1.000 (0.000) |
| S6 | 2-state HMM | 2-state HSMM | Duration-aware | 2 | 2 | 300 | 250 | $\log(20)$ | 0.000 (0.000);<br>1.000 (0.000) |
| S7 | 2-state HMM | 2-state (localized fail.) | State-localized | 2 | 2 | 300 | 300 | $\log(20)$ | 0.063 (0.014);<br>0.900 (0.017) |
| S8 | 2-state HMM | 2-state (change point) | Switching mix | 2 | 2 | 300 | 250 | $\log(20)$ | 0.036 (0.012);<br>1.000 (0.000) |

The calibration entries in Table 1 show false-signal behavior compatible with the anytime-valid threshold: S1 crosses at 0.030, the null runs in S6 and S8 cross at 0.000 and 0.036, and S7 crosses at 0.063, slightly above 0.05 but within Monte Carlo error (SE 0.014).

The alternative rows in Table 1 show targeted sensitivity under controlled misspecification. S3 detects a missing behavioural state with crossing rate 0.997; S4, S5 and S6 detect their respective targeted failures in all replicates; and S7 reaches 0.900 for a localized failure. These near-perfect rates reflect deliberately targeted alternatives, not broad power guarantees.

The main lesson from Table 1 is that e-diagnostics are targeted predictive comparisons. They grow when the diagnostic density predicts better than the fitted null and remain calibrated when the fitted generator is the validation law. Power may be lower for highly overlapping state-dependent distributions, small angular misspecification, moderate copula dependence, near-geometric dwell-time distributions or alternatives estimated from limited training data.

Additional scenarios are reported in Online Supplement S5, and Online Supplement S6 gives a retrospective pseudo-residual comparison (Zucchini et al., 2016).

## 7 Elk movement application

The purpose of this application is not to draw definitive biological conclusions about elk behaviour, but to illustrate how sequential predictive e-diagnostics identify individual-specific and state-localized predictive failures in a standard movement-HMM workflow.

We used `moveHMM::elk_data` (Michelot et al., 2016), associated with Morales et al. (2004). The dataset contains four individuals and 735 projected locations with distance to water. We computed step lengths and turning angles with `moveHMM::prepData`. Under the LOAO protocol, each individual was held out while candidate models were trained on the other three. Figure 1 displays the trajectories.

**Figure 1:**
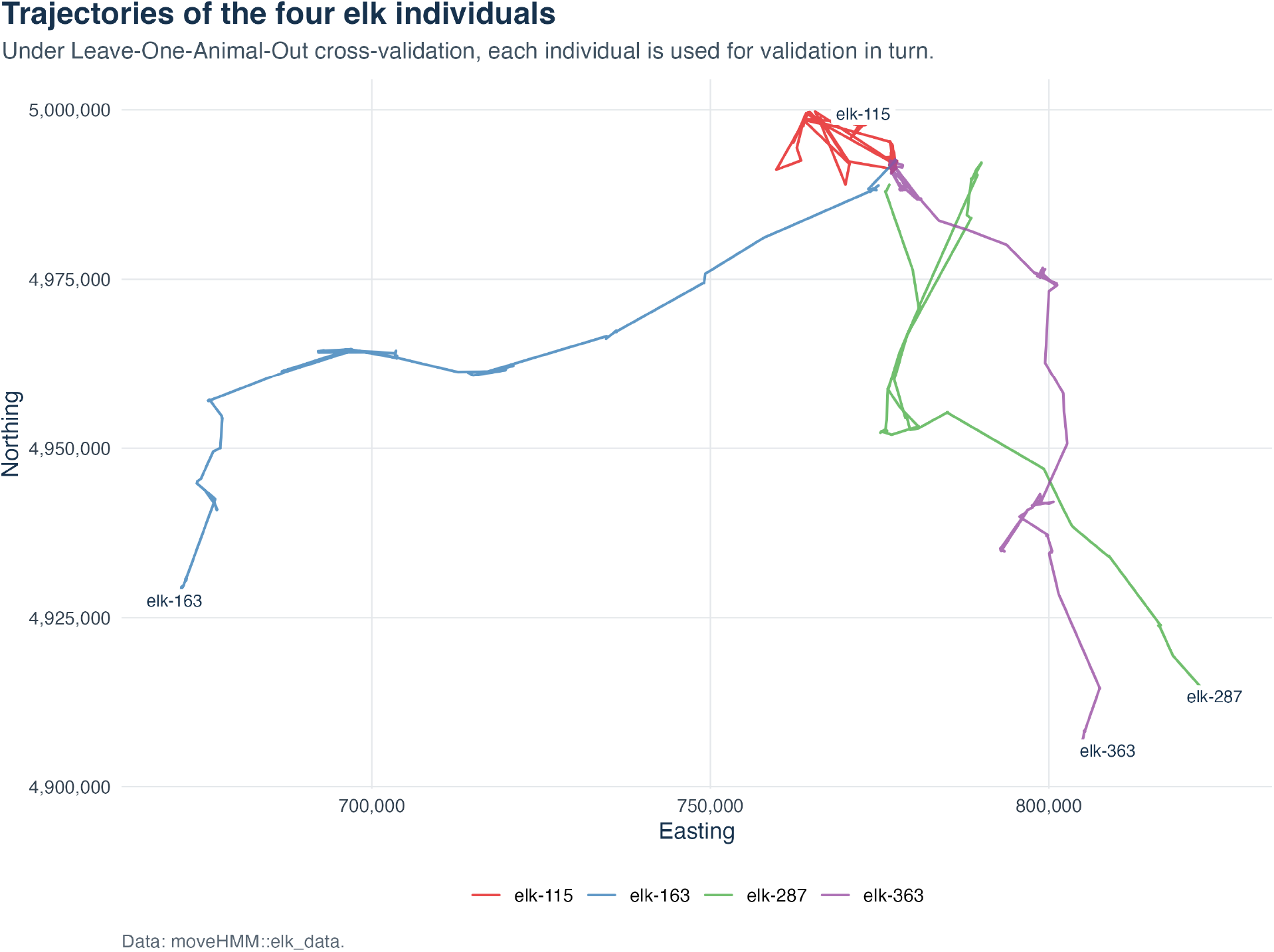
Trajectories of the four elk individuals. Under the LOAO protocol, each trajectory is used for validation in turn.

For each fold, we fitted HMMs with *K* = 2, 3, 4 states, Gamma step lengths, von Mises turning angles and no transition covariates. Fits used `moveHMM::fitHMM` from five deterministic starts; states were ordered by mean step length; and the filter was restarted for each validation individual. AIC and BIC values are in Online Supplement S1. A pre-specified majority-BIC rule fixed the null to *K* = 3 and the diagnostic alternative to *K* = 4.

We evaluated four pre-specified diagnostics: a full-density *K* + 1 diagnostic, a diffuse-angle diagnostic, their equal-weight mixture, and a state-localized *K* +1 diagnostic using the filtered probability of the long-step state. Figure 2 shows the cumulative paths, and Table 2 gives final individual and population-average e-values (Proposition 5).

**Table 2:** LOAO predictive e-diagnostics for the four elk individuals and the population-average cross-fitted e-values. The threshold for *α* = 0.05 is log(20) = 2.996. Crossing times and maximum log e-values are given in Online Supplement S1.

| Validation individual | Diagnostic | Signal | Final log $e$ |
| --- | --- | --- | --- |
| elk-115 | $K = 3$ vs $K = 4$ | Yes | 5.413 |
| elk-115 | Diffuse angle | No | 0.154 |
| elk-115 | Fixed mixture | Yes | 10.981 |
| elk-115 | Localized $K + 1$ | Yes | 5.605 |
| elk-163 | $K = 3$ vs $K = 4$ | No | -12.216 |
| elk-163 | Diffuse angle | Yes | 2.663 |
| elk-163 | Fixed mixture | No | -1.244 |
| elk-163 | Localized $K + 1$ | No | -1.725 |
| elk-287 | $K = 3$ vs $K = 4$ | No | 1.056 |
| elk-287 | Diffuse angle | No | 1.801 |
| elk-287 | Fixed mixture | No | 2.592 |
| elk-287 | Localized $K + 1$ | No | 0.513 |
| elk-363 | $K = 3$ vs $K = 4$ | Yes | 7.246 |
| elk-363 | Diffuse angle | No | -0.761 |
| elk-363 | Fixed mixture | Yes | 9.174 |
| elk-363 | Localized $K + 1$ | Yes | 4.734 |
| Average | $K = 3$ vs $K = 4$ | Yes | 6.009 |
| Average | Diffuse angle | No | 1.706 |
| Average | Fixed mixture | Yes | 9.747 |
| Average | Localized $K + 1$ | Yes | 4.573 |

**Figure 2:**
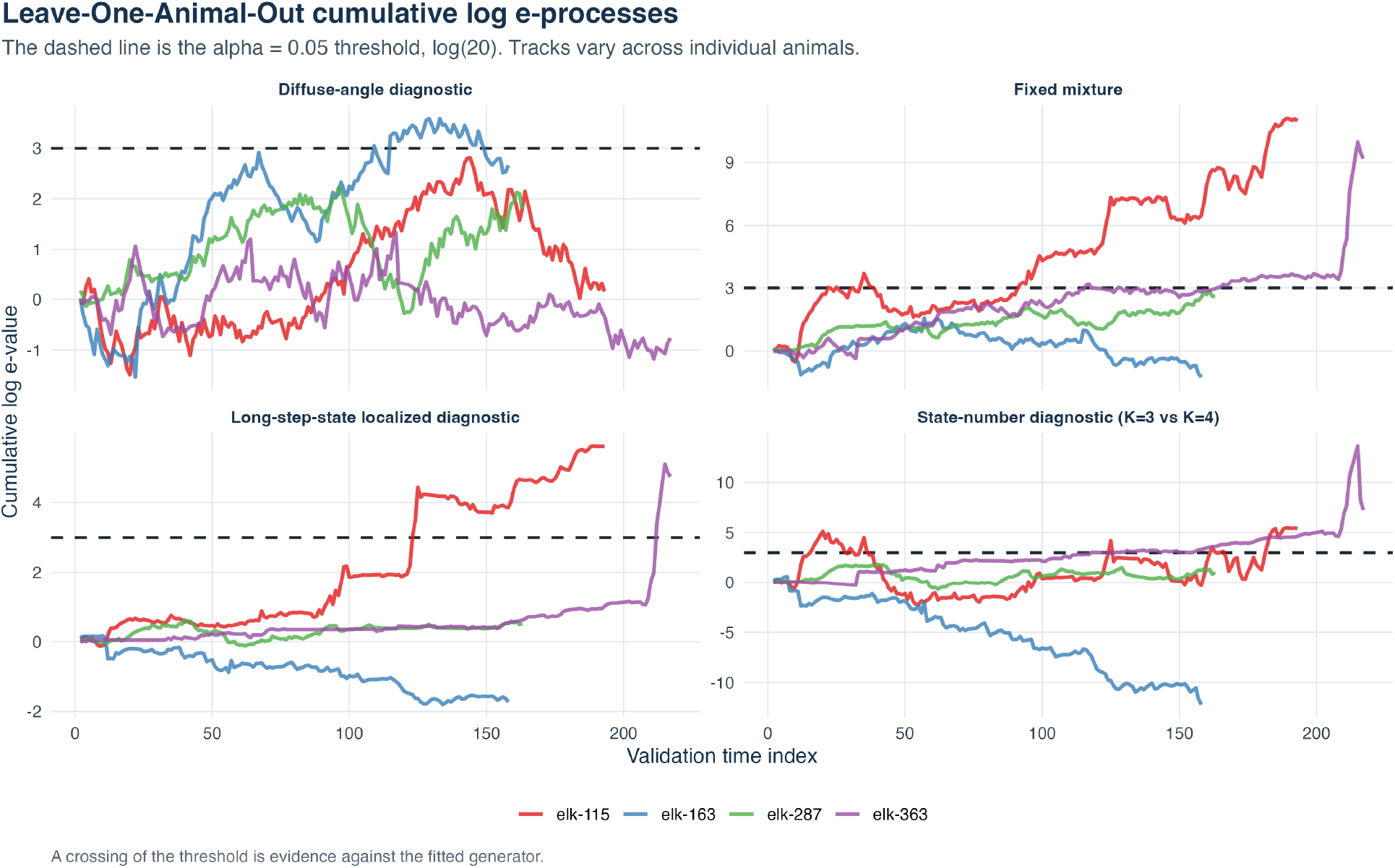
Cumulative log e-processes for the four validation elk individuals across the four diagnostics. The dashed horizontal line is the *α* = 0.05 threshold, log(20) = 2.996.

The diagnostics vary across individuals. For `elk-115` and `elk-363`, the *K* + 1 state diagnostic and fixed mixture cross log(20) = 2.996; the state-localized diagnostic also crosses for these two individuals. For `elk-163`, the diffuse-angle diagnostic gives the main signal, whereas `elk-287` shows no clear crossing.

For joint confirmatory statements across *J* = 4 diagnostics, a Bonferroni union-bound threshold is log(*J/α*) = log(80) ≈ 4.382. Under this conservative threshold, the number-of-states diagnostic and fixed mixture still cross for `elk-115` and `elk-363`, and the state-localized diagnostic still crosses for `elk-115`; the diffuse-angle diagnostic for elk-163 no longer crosses.

Population-average pooling, 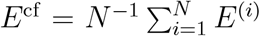, is valid when the fold-specific validation laws are embedded in a common joint null under which each *E*^(*i*)^ has expectation at most one. Because LOAO training sets overlap, this is a cross-predictive diagnostic rather than an automatic exact global test. The average can also be dominated by large individual e-values, so individual paths remain essential.

At the population level, Table 2 exceeds the terminal threshold for the number-of-states, fixed-mixture and localized diagnostics, but not for the diffuse-angle diagnostic. In this illustrative dataset, lack of fit is concentrated in some held-out individuals and in the number-of-states and long-step-localized diagnostics. Figure 3 shows the predictable state weights and localized e-process for `elk-115`.

**Figure 3:**
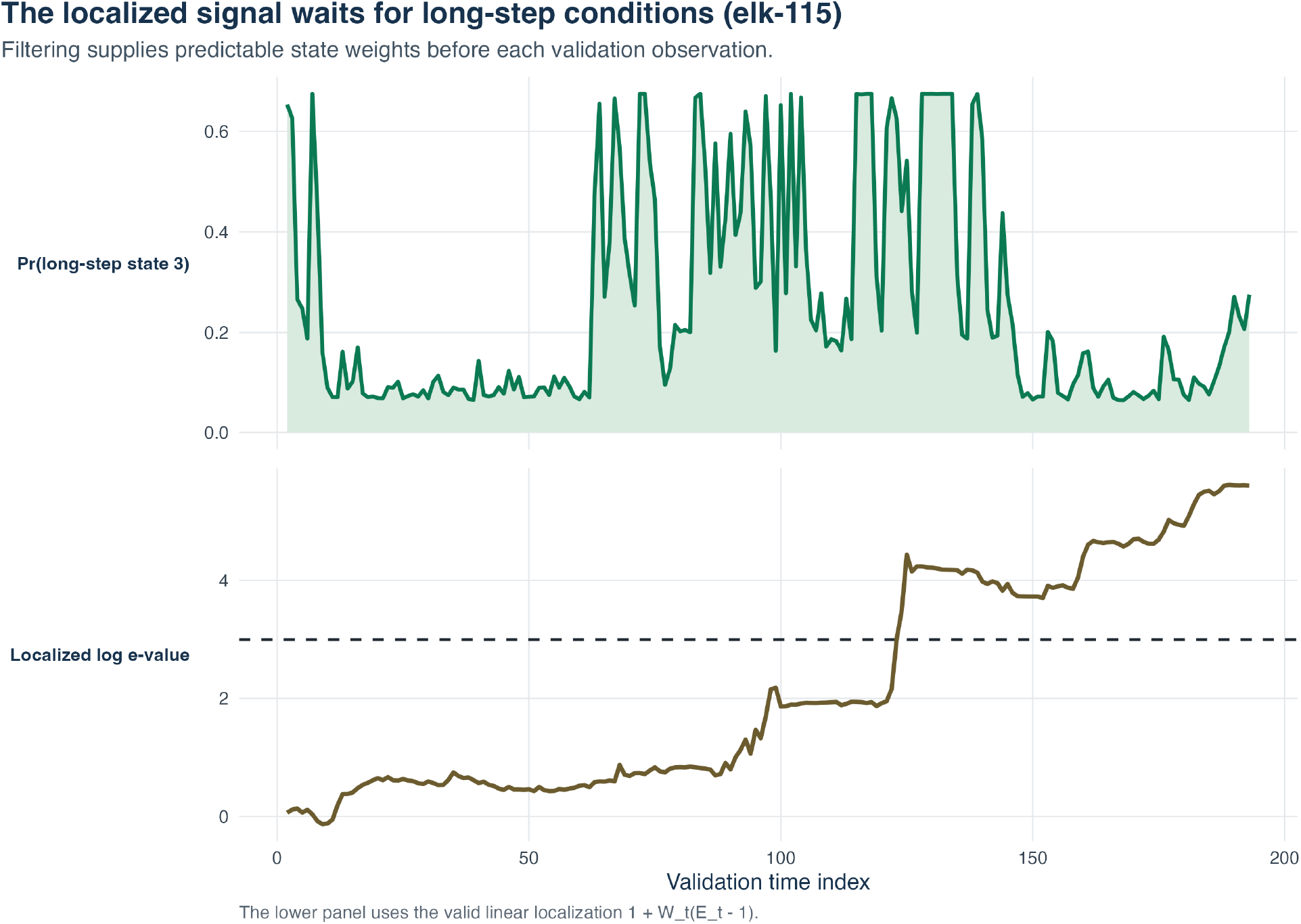
Predictable state-localized diagnostic for validation individual `elk-115`. The upper panel shows the filtered predictive probability assigned to the long-step state before each validation observation. The lower panel shows the corresponding localized cumulative log e-process.

Thus the LOAO diagnostics suggest individual-specific predictive lack of fit and, with the caution above, provide a population-level terminal summary.

## 8 Discussion

This paper develops a predictive framework for checking hidden Markov models of animal movement. A fitted HMM is treated as a sequential generator of held-out data: it supplies the observable one-step predictive density, a diagnostic alternative supplies a predictable competitor, and the cumulative ratio is an e-process under the fixed fitted-generator null.

The central technical point is that the denominator must be the marginal observable predictive density obtained by filtering over latent states. Conditioning instead on a Viterbi or locally decoded behavioural state can invalidate the e-value, because decoded states are data-dependent reconstructions and may use the current or future observations. Thus decoded states remain useful for interpretation after a signal is found, but not as denominators for predictive validity.

The movement-oriented extensions include predictable mixtures and switching, localization over seasons, habitats, individuals or likely states, feature-level circular-linear diagnostics, and blockwise checks of displacement, straightness, residence, barrier crossing or space use. These tools reuse state-dependent densities and filtered probabilities already computed in standard HMM workflows.

The simulations illustrate calibration and targeted sensitivity. Under the fitted-generator null, empirical crossing rates were compatible with the anytime-valid guarantee; under controlled alternatives, evidence accumulated for missing states, angular misspecification, residual dependence and non-geometric dwell times. Power depends on how well the diagnostic alternative predicts the validation sequence.

The elk application illustrates the supported level of applied interpretation. The LOAO analysis suggests individual-specific predictive lack of fit in the fitted three-state generator, with stronger *K* + 1 and mixture signals for `elk-115` and `elk-363`, a more angular signal for `elk-163`, and no clear signal for `elk-287`. The population-average e-value is useful as a cross-predictive summary, but is not an automatic exact global test unless the fold-specific validation laws are embedded in an explicit joint null.

The main limitation is the same feature that gives the method its clarity. Exact validity is strongest when the fitted null, preprocessing and diagnostic alternatives are fixed before validation. The results do not fully account for estimated-parameter uncertainty and do not make same-sample exploratory workflows confirmatory. Composite-null envelopes can give uniform validity, but conservatively, and richer alternatives may add computational burden for covariate-dependent HMMs, HSMMs or blockwise spatial summaries.

Future work should sharpen treatment of parameter uncertainty, develop composite-null constructions beyond finite or compact families, and study diagnostics estimated from limited training data. Extensions to environmental covariates, step-selection and HMM-SSF models, hidden semi-Markov models, spatial block diagnostics and dedicated software are natural next steps. The framework complements existing diagnostics by making predictive failure explicit in time, location and direction, while leaving ecological interpretation to the analyst and scientific context.

## Supporting information

supplementary materiel

## Data Availability Statement

The elk movement data analysed in this paper are publicly available in the R package `moveHMM` (Michelot et al., 2016) and were originally collected for the movement study of Morales et al. (2004). Code, simulation parameters, random seeds, and documentation needed to reproduce the tables and figures are available at https://github.com/AurelienNicosiaULaval/evalue-HMM. Package documentation is available at https://aureliennicosiaulaval.github.io/evalue-HMM/.

