## supplementary materiel for "Sequential predictive e-diagnostics for hidden Markov models of animal movement"

#### S1 Supplementary notation, tables, and figures

This section collects supporting tables and figures referenced from the main article, namely the notation summary, the statistical-target summary, the elk model-selection metrics, the detailed elk diagnostic table and the population-level summary of the elk diagnostics.

| Notation | Meaning |
|---|---|
| $Y_t$ | Observed movement variable at time $t$, often $(L_t, \Theta_t)$ |
| $L_t$ | Step length at time $t$ |
| $\Theta_t$ | Turning angle at time $t$ |
| $S_t$ | Latent behavioural state at time $t$ |
| $K$ | Number of states in the null HMM |
| $a_{1\|0}(s)$ | Initial predictive probability for state $s$ |
| $\mathcal{H}$ | Training information fixed before validation |
| $\mathcal{F}_t$ | Observable validation filtration, $\sigma(\mathcal{H}, Y_1, \ldots, Y_t)$ |
| $M_0$ | Null HMM or fitted null generator |
| $p_0(y_t \mid \mathcal{F}_{t-1})$ | Observable predictive density under $M_0$ |
| $q_t(y_t \mid \mathcal{F}_{t-1})$ | Diagnostic predictive density or subdensity at time $t$ |
| $E_t$ | Predictive e-value increment at time $t$ |
| $E_{1:t}$ | Cumulative product of increments up to time $t$ |
| $\alpha$ | Anytime-valid monitoring level |

Table S1: Notation used throughout the theoretical setup of the main article.

#### S2 Additional theoretical results

This section collects results referenced from the main article: an anytime p-value formulation induced by the e-process, a counterexample showing why a decoded-state denominator can violate e-validity, a conservative composite-null envelope (which underlies scenario S12), and a log-score characterization of e-process growth under misspecification.

| Target | Validity claim | Practical interpretation |
|---|---|---|
| Fixed fitted generator | Exact conditional e-process | "Does this fitted HMM generate the validation trajectory?" |
| True parametric HMM with estimated parameters | Not exact without adjustment | Requires parameter uncertainty treatment |
| Composite family | Uniform e-process via envelope or other safe construction | Conservative but genuine family-wise null |

Table S2: Distinct statistical targets for movement model validation. The main article focuses primarily on the fixed fitted generator, treating the diagnostic as a check of the specific predictive model resulting from training.

| Fold (Validation Indiv.) | States ($K$) | Negative Log-Likelihood | Parameters | AIC | BIC |
|---|---|---|---|---|---|
| **elk-115** | 2 | 5190.649 | 13 | 10407.298 | 10462.919 |
| (Training $n = 533$) | 3 | 4994.368 | 23 | 10034.736 | 10133.142$^\star$ |
| | 4 | 4976.707 | 35 | 10023.414 | 10173.162 |
| **elk-163** | 2 | 5442.706 | 13 | 10911.412 | 10967.860 |
| (Training $n = 568$) | 3 | 5378.781 | 23 | 10803.562 | 10903.430 |
| | 4 | 5335.920 | 35 | 10741.841 | 10893.815$^\star$ |
| **elk-287** | 2 | 5481.246 | 13 | 10988.493 | 11044.826 |
| (Training $n = 563$) | 3 | 5332.742 | 23 | 10711.484 | 10811.149$^\star$ |
| | 4 | 5297.755 | 35 | 10665.510 | 10817.175 |
| **elk-363** | 2 | 4999.619 | 11 | 10021.237 | 10067.837 |
| (Training $n = 511$) | 3 | 4830.232 | 20 | 9700.463 | 9785.190$^\star$ |
| | 4 | 4813.181 | 31 | 9688.363 | 9819.690 |

Table S3: Model selection metrics for fitted HMMs across leave-one-animal-out (LOAO) training sets. The $^\star$ symbol denotes the model selected by BIC for each fold.

#### S2.1 Anytime p-value induced by an e-process

**Corollary 1** (Anytime p-value induced by an e-process)**.** *Let $(E_{1:t})_{0\le t\le T}$ be the predictive e-process of the main article. Define*

$$p_t^{\text{any}} = \min\left\{1, \frac{1}{\sup_{0\le u\le t} E_{1:u}}\right\}. \quad (1)$$

*Then $p_t^{\text{any}}$ is anytime-valid in the sense that, under the null generator,*

$$\mathbb{P}_0\left(\exists\, 0 \le t \le T : p_t^{\text{any}} \le \alpha\right) \le \alpha. \quad (2)$$

*Proof.* The event $\{\exists\, 0 \le t \le T : p_t^{\text{any}} \le \alpha\}$ is the same as $\{\sup_{0\le t\le T} E_{1:t} \ge 1/\alpha\}$, which is controlled by the optional-stopping and anytime-monitoring corollary in the main article. □

*Remark* 1 (Support convention)*.* When $E_t$ is set to zero on $\{p_0 = 0\}$, any diagnostic mass assigned outside the null support is truncated. This preserves e-process validity under the null validation law, but it does not use discrepancies on events that the null assigns probability zero.

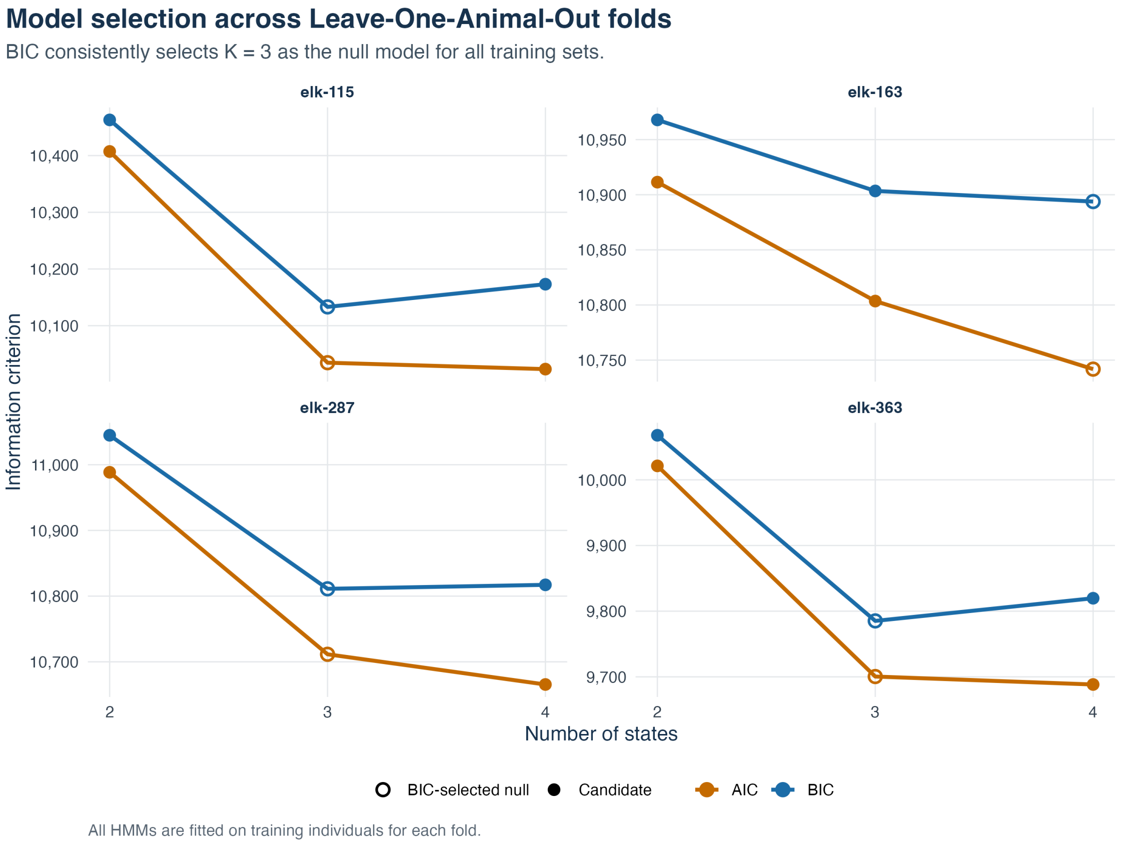


Figure S1: AIC and BIC for the fitted HMMs across the four elk training folds. BIC selects $K = 3$ for three folds and $K = 4$ for one fold.

#### S2.2 A state-conditioned denominator can violate e-validity

**Example 1** (A state-conditioned denominator can violate e-validity)**.** Consider a two-state model at a single validation time with predictive state probabilities 1/2 and 1/2. Let

$$f_1(y) = \phi(y; -m, 1), \qquad f_2(y) = \phi(y; m, 1),$$

where $\phi(\cdot; \mu, 1)$ is the $N(\mu, 1)$ density and $m > 0$. The observable null density is

$$p_0(y) = \frac{1}{2} f_1(y) + \frac{1}{2} f_2(y).$$

Suppose one incorrectly uses $f_1(y)$ as the denominator because a decoded or assigned state is set to state 1, and suppose the diagnostic numerator is $q(y) = f_2(y)$. The expected ratio under the true observable null is

$$\mathbb{E}_0 \left\{ \frac{f_2(Y)}{f_1(Y)} \right\} = \int \frac{f_2(y)}{f_1(y)} \left\{ \frac{1}{2} f_1(y) + \frac{1}{2} f_2(y) \right\} \mathrm{d}y \tag{3}$$

$$= \frac{1}{2} + \frac{1}{2} \int \frac{f_2(y)^2}{f_1(y)} \mathrm{d}y \tag{4}$$

$$= \frac{1}{2} + \frac{1}{2} \exp(4m^2), \tag{5}$$

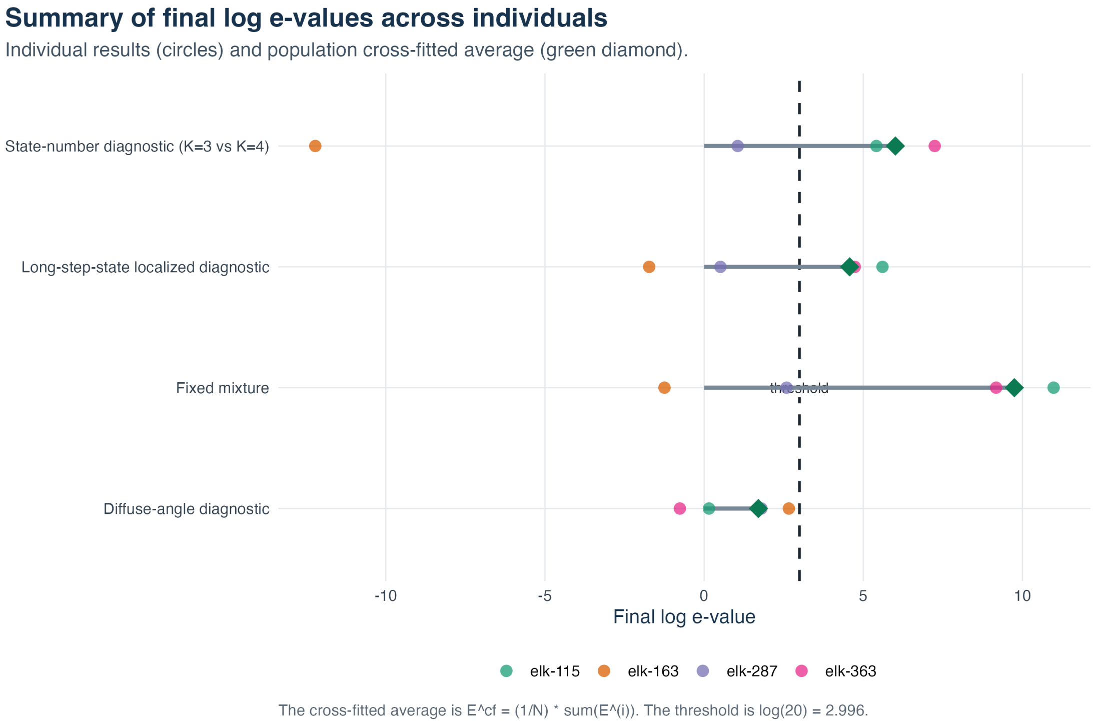


Figure S2: Summary of final log e-values across validation individuals (circles) and the population-average cross-fitted e-value (diamond marker). The dashed line represents the $\alpha = 0.05$ threshold, $\log(20) = 2.996$.

which is larger than 1. Thus the ratio is not an e-value under the observable null. The problem is not the use of likelihood ratios; the problem is the use of the wrong denominator. The denominator must always be the filtered marginal predictive density $p_0(Y_t \mid \mathcal{F}_{t-1})$.

#### S2.3 Conservative envelope for composite null families

**Proposition 2** (Composite-null envelope). *Let $\{p_\theta(y \mid \mathcal{F}_{t-1}) : \theta \in \Theta_0\}$ be a family of null predictive densities. Assume that the pointwise supremum*

$$p_t^{\text{env}}(y \mid \mathcal{F}_{t-1}) = \sup_{\theta \in \Theta_0} p_\theta(y \mid \mathcal{F}_{t-1}) \tag{6}$$

*is measurable, positive wherever any $p_\theta$ is positive, and finite on the relevant support. This is typically satisfied if $\Theta_0$ is a finite family, or by using an essential supremum over a well-controlled parametric family. Let $q_t$ be a predictable diagnostic subdensity. Define*

$$\overline{E}_t = \frac{q_t(Y_t \mid \mathcal{F}_{t-1})}{p_t^{\text{env}}(Y_t \mid \mathcal{F}_{t-1})}. \tag{7}$$

*Then for every $\theta \in \Theta_0$,*

$$\mathbb{E}_\theta(\overline{E}_t \mid \mathcal{F}_{t-1}) \le 1, \tag{8}$$

*and $\prod_{u \le t} \overline{E}_u$ is an e-process uniformly valid over the composite null family.*

| Validation individual | Diagnostic | Crossing time | Max $\log e$ |
|---|---|---|---|
| `elk-115` | $K = 3$ vs $K = 4$ | 16 | 5.459 |
| `elk-115` | Diffuse angle | NA | 2.811 |
| `elk-115` | Fixed mixture | 22 | 11.089 |
| `elk-115` | Localized $K + 1$ | 123 | 5.618 |
| `elk-163` | $K = 3$ vs $K = 4$ | NA | 0.610 |
| `elk-163` | Diffuse angle | 109 | 3.585 |
| `elk-163` | Fixed mixture | NA | 1.556 |
| `elk-163` | Localized $K + 1$ | NA | 0.165 |
| `elk-287` | $K = 3$ vs $K = 4$ | NA | 1.854 |
| `elk-287` | Diffuse angle | NA | 2.220 |
| `elk-287` | Fixed mixture | NA | 2.836 |
| `elk-287` | Localized $K + 1$ | NA | 0.619 |
| `elk-363` | $K = 3$ vs $K = 4$ | 118 | 13.627 |
| `elk-363` | Diffuse angle | NA | 1.337 |
| `elk-363` | Fixed mixture | 116 | 9.973 |
| `elk-363` | Localized $K + 1$ | 212 | 5.095 |
| Average | $K = 3$ vs $K = 4$ | NA | 6.009 |
| Average | Diffuse angle | NA | 1.706 |
| Average | Fixed mixture | NA | 9.747 |
| Average | Localized $K + 1$ | NA | 4.573 |

Table S4: Crossing times and maximum log e-values for the elk diagnostics summarized in Table 2 of the main article. For population-average rows, the maximum equals the terminal value by definition.

*Proof.* For each fixed $\theta \in \Theta_0$, $p_\theta(y \mid \mathcal{F}_{t-1}) \le p_t^{\text{env}}(y \mid \mathcal{F}_{t-1})$ pointwise. Hence

$$\mathbb{E}_\theta(\overline{E}_t \mid \mathcal{F}_{t-1}) = \int \frac{q_t(y \mid \mathcal{F}_{t-1})}{p_t^{\text{env}}(y \mid \mathcal{F}_{t-1})} p_\theta(y \mid \mathcal{F}_{t-1}) \mu_t(\mathrm{d}y) \tag{9}$$

$$\le \int q_t(y \mid \mathcal{F}_{t-1}) \mu_t(\mathrm{d}y) \tag{10}$$

$$\le 1. \tag{11}$$

The product result follows from the supermartingale argument. □

*Remark* 2 (Cost and limits of envelope validity). The envelope denominator is usually not a density and may be much larger than any single fitted predictive density. The resulting e-process can be conservative and computationally difficult for HMMs. Moreover, continuous envelopes over unrestricted Gamma or von Mises families may be infinite, unstable, or computationally prohibitive. The envelope construction is therefore primarily practical for finite or compactly restricted parameter families, rather than general HMM parameter spaces.

#### S2.4 Behaviour under misspecification

**Proposition 3** (Conditional log-score decomposition)**.** *Let $p_\star(\cdot \mid \mathcal{F}_{t-1})$ be the true conditional density of $Y_t$ under a data-generating process $P_\star$. Let $q_t$ and $p_0$ be predictive densities. If the relevant integrals are finite, then*

$$\mathbb{E}_\star\left[\log \frac{q_t(Y_t \mid \mathcal{F}_{t-1})}{p_0(Y_t \mid \mathcal{F}_{t-1})} \middle| \mathcal{F}_{t-1}\right] = \mathrm{KL}\{p_\star(\cdot \mid \mathcal{F}_{t-1}), p_0(\cdot \mid \mathcal{F}_{t-1})\} \tag{12}$$

$$- \mathrm{KL}\{p_\star(\cdot \mid \mathcal{F}_{t-1}), q_t(\cdot \mid \mathcal{F}_{t-1})\}. \tag{13}$$

*Thus the expected local log-increment is positive exactly when the diagnostic alternative is closer than the null in conditional Kullback-Leibler risk.*

*Proof.* By definition,

$$\mathbb{E}_\star\left[\log\frac{q_t(Y_t \mid \mathcal{F}_{t-1})}{p_0(Y_t \mid \mathcal{F}_{t-1})}\middle|\mathcal{F}_{t-1}\right] = \int p_\star(y \mid \mathcal{F}_{t-1})\log\frac{q_t(y \mid \mathcal{F}_{t-1})}{p_0(y \mid \mathcal{F}_{t-1})}\mu_t(\mathrm{d}y) \tag{14}$$

$$= \int p_\star \log\frac{p_\star}{p_0}\mu_t(\mathrm{d}y) - \int p_\star \log\frac{p_\star}{q_t}\mu_t(\mathrm{d}y), \tag{15}$$

which is the displayed identity. □

**Proposition 4** (Asymptotic growth under positive drift)**.** *Let $X_t = \log E_t$ and suppose under $P_\star$ that $\delta_t = \mathbb{E}_\star(X_t \mid \mathcal{F}_{t-1})$ exists. Assume the martingale differences $X_t - \delta_t$ satisfy a strong law, for example*

$$\frac{1}{T}\sum_{t=1}^{T}\{X_t - \delta_t\} \to 0 \qquad P_\star\text{-a.s.} \tag{16}$$

*If $\frac{1}{T}\sum_{t=1}^{T}\delta_t \to \delta > 0$, then*

$$\frac{1}{T}\log E_{1:T} \to \delta \qquad P_\star\text{-a.s.} \tag{17}$$

*In particular, the e-process grows exponentially at asymptotic rate $\delta$.*

*Proof.* Using $\log E_{1:T} = \sum_{t=1}^{T} X_t$,

$$\frac{1}{T}\log E_{1:T} = \frac{1}{T}\sum_{t=1}^{T}\delta_t + \frac{1}{T}\sum_{t=1}^{T}(X_t - \delta_t). \tag{18}$$

The first term converges to $\delta$ by assumption and the second to zero by the assumed martingale strong law. □

*Remark* 3 (Power is diagnostic-specific)*.* A large e-process is evidence against the null relative to the alternative used to construct the diagnostic. If the null HMM has an ecologically important defect but the alternative does not predict that defect better, the e-process may not grow. This is a feature rather than a failure: the method localizes model criticism through explicit diagnostic bets.

### S3 Predictable mixtures, localization, and feature-level diagnostics

This section collects methodological results referenced from the main article: predictable mixtures and switching, localization and tempering, feature-level and blockwise diagnostics, the movement-specific diagnostic catalogue, and the recommended implementation workflow.

#### S3.1 Predictable diagnostic selection, mixtures and parallel diagnostics

In practice, the analyst may not want to commit to a single diagnostic alternative. One may want to compare a $K+1$ state HMM, an angular alternative, a copula alternative and a duration alternative. E-values are convenient here, but care is needed. Predictable switching and mixtures are valid. Products of parallel diagnostics computed on the same validation observations are not automatically valid.

**Proposition 5** (Predictable mixtures of diagnostic alternatives). *Let $q_{j,t}(y \mid \mathcal{F}_{t-1})$, $j = 1, \ldots, J$, be predictable diagnostic subdensities satisfying*

$$\int q_{j,t}(y \mid \mathcal{F}_{t-1}) \mu_t(\mathrm{d}y) \leq 1.$$

*Let $w_{j,t} \geq 0$ be predictable weights with $\sum_{j=1}^{J} w_{j,t} \leq 1$. Define*

$$q_t^{\text{mix}}(y \mid \mathcal{F}_{t-1}) = \sum_{j=1}^{J} w_{j,t} q_{j,t}(y \mid \mathcal{F}_{t-1}) \tag{19}$$

*and*

$$E_t^{\text{mix}} = \frac{q_t^{\text{mix}}(Y_t \mid \mathcal{F}_{t-1})}{p_0(Y_t \mid \mathcal{F}_{t-1})}. \tag{20}$$

*Then $E_t^{\text{mix}}$ is a conditional e-value increment, and $\prod_{u \leq t} E_u^{\text{mix}}$ is an e-process under $M_0$.*

*Proof.* The mixture $q_t^{\text{mix}}$ is predictable and integrates to at most $\sum_j w_{j,t} \leq 1$. The result follows directly from Theorem 1 of the main article. □

**Corollary 6** (Predictable switching). *Let $J_t \in \{1, \ldots, J\}$ be $\mathcal{F}_{t-1}$-measurable. Then*

$$E_t^{\text{switch}} = \frac{q_{J_t,t}(Y_t \mid \mathcal{F}_{t-1})}{p_0(Y_t \mid \mathcal{F}_{t-1})} \tag{21}$$

*is a conditional e-value increment. Thus the analyst may switch among a pre-specified menu of diagnostic alternatives using only information available before the next validation observation.*

*Proof.* This is Proposition 5 with $w_{j,t} = \mathbf{1}\{J_t = j\}$. □

**Proposition 7** (Parallel diagnostics on the same validation data). *Suppose $E^{(1)}, \ldots, E^{(J)}$ are e-values under $M_0$ computed from the same validation experiment, so that $\mathbb{E}_0 E^{(j)} \leq 1$ for each $j$. If $w_j \geq 0$ and $\sum_j w_j \leq 1$, then*

$$E^{\text{avg}} = \sum_{j=1}^{J} w_j E^{(j)} \tag{22}$$

*is also an e-value under $M_0$. Consequently,*

$$\mathbb{P}_0 \left( \max_{1 \leq j \leq J} w_j E^{(j)} \geq \frac{1}{\alpha} \right) \leq \alpha. \tag{23}$$

*In contrast, the product $\prod_{j=1}^{J} E^{(j)}$ is not generally an e-value when the diagnostics are computed in parallel on the same observations.*

*Proof.* Linearity of expectation gives

$$\mathbb{E}_0 E^{\text{avg}} = \sum_j w_j \mathbb{E}_0 E^{(j)} \le \sum_j w_j \le 1.$$

If $\max_j w_j E^{(j)} \ge 1/\alpha$, then $E^{\text{avg}} \ge 1/\alpha$, so Markov's inequality applied to the e-value $E^{\text{avg}}$ gives the displayed bound. The final statement is a warning: without additional conditional independence or a sequential construction using fresh information, products of e-values need not have expectation at most one. □

*Remark* 4 (Recommended reporting for several diagnostics). For a diagnostic menu evaluated on the same validation sequence, report the individual e-processes and either use a weighted average e-value or pre-assigned weighted thresholds. A product of diagnostic e-processes is valid only when the factors are accumulated as a sequential process with each new factor conditionally valid given all previous diagnostic outcomes and using the appropriate next unit of information.

#### S3.2 Localization, state weighting and tempering

A global e-process answers whether a diagnostic alternative accumulates evidence against the null generator. Movement ecologists usually also want to know where the failure occurs: during migration, in a given habitat, for one individual, or during time periods in which the fitted model predicts a particular behavioural state. The following result gives a valid way to localize evidence when the localization weights are predictable.

**Proposition 8** (Predictable weighted localization). *Let $E_t$ be any conditional e-value increment satisfying $\mathbb{E}_0(E_t \mid \mathcal{F}_{t-1}) \le 1$. Let $W_t$ be any $\mathcal{F}_{t-1}$-measurable weight with $0 \le W_t \le 1$. Define*

$$E_t^{(W)} = 1 + W_t(E_t - 1). \tag{24}$$

*Then $E_t^{(W)} \ge 0$ and*

$$\mathbb{E}_0(E_t^{(W)} \mid \mathcal{F}_{t-1}) \le 1. \tag{25}$$

*Consequently, $\prod_{u \le t} E_u^{(W)}$ is an e-process under $M_0$.*

*Proof.* Since $E_t \ge 0$ and $0 \le W_t \le 1$, $E_t^{(W)} = (1 - W_t) + W_t E_t \ge 0$. Also,

$$\mathbb{E}_0(E_t^{(W)} \mid \mathcal{F}_{t-1}) = 1 + W_t\{\mathbb{E}_0(E_t \mid \mathcal{F}_{t-1}) - 1\} \tag{26}$$

$$\le 1. \tag{27}$$

The product result follows from the same supermartingale argument as in Theorem 1 of the main article. □

**Corollary 9** (Segment-specific e-processes). *If $B_t$ is a predictable indicator of a biologically meaningful segment, such as a known season, a habitat class known before observing the movement response, or a held-out individual, then*

$$E_t^{(B)} = \begin{cases} E_t, & B_t = 1, \\ 1, & B_t = 0, \end{cases} \tag{28}$$

*yields a valid segment-specific e-process. Thus evidence can be accumulated only over a pre-specified or predictably selected subset of times.*

*Proof.* This is Proposition 8 with $W_t = B_t \in \{0, 1\}$. $\square$

**Corollary 10** (Soft state-localized e-processes)**.** *For a fitted HMM, set*

$$W_t^{(s)} = a_{t|t-1}(s) = \mathbb{P}_0(S_t = s \mid \mathcal{F}_{t-1}). \tag{29}$$

*Then*

$$E_t^{(s)} = 1 + a_{t|t-1}(s)(E_t - 1) \tag{30}$$

*defines a valid e-process localized to times at which state $s$ is predicted to be likely under the null model.*

*Proof.* The filtered predictive probability $a_{t|t-1}(s)$ is $\mathcal{F}_{t-1}$-measurable and lies in $[0, 1]$, so Proposition 8 applies. $\square$

**Proposition 11** (Power-tempered and linearly tempered e-increments)**.** *Let $E_t$ be a conditional e-value increment. If $\lambda_t \in [0, 1]$ is predictable, then both*

$$\widetilde{E}_t^{\text{pow}} = E_t^{\lambda_t} \tag{31}$$

*(with the convention $0^0 = 1$) and*

$$\widetilde{E}_t^{\text{lin}} = 1 + \lambda_t(E_t - 1) \tag{32}$$

*are conditional e-value increments. Their products are e-processes.*

*Proof.* The linear case follows from Proposition 8 with $W_t = \lambda_t$. For the power case, Jensen's inequality applied to the concave function $x \mapsto x^{\lambda_t}$ gives

$$\mathbb{E}_0(E_t^{\lambda_t} \mid \mathcal{F}_{t-1}) \le \{\mathbb{E}_0(E_t \mid \mathcal{F}_{t-1})\}^{\lambda_t} \le 1.$$

$\square$

*Remark* 5 (Interpretation of localized evidence)*.* The process in Corollary 10 is not the same as conditioning on the event that the true latent state equals $s$. It is a valid soft localization based on the null model's predictive probability of state $s$. Smoothed state probabilities using future observations can be useful for visualization, but they are not predictable and therefore do not by themselves preserve the e-process guarantee.

#### S3.3 Feature-level and blockwise e-diagnostics

The previous sections use full predictive densities for $Y_t$. In movement applications, the diagnostically relevant object may be a lower-dimensional feature, such as a circular residual, a transformed step length, a copula residual, a barrier-crossing indicator or a blockwise spatial statistic. E-values can be constructed directly from such features if their null conditional law is known.

**Theorem 12** (Feature-level predictive e-values)**.** *Let $Z_t = \psi_t(Y_t, \mathcal{F}_{t-1})$ be a diagnostic feature whose transformation rule $\psi_t$ is predictable. Let $g_{0,t}(z \mid \mathcal{F}_{t-1})$ be the conditional density of $Z_t$ under $M_0$, and let $g_{1,t}(z \mid \mathcal{F}_{t-1})$ be a predictable diagnostic subdensity satisfying $\int g_{1,t}(z \mid \mathcal{F}_{t-1})\nu_t(\mathrm{d}z) \leq 1$. Define*

$$R_t = \frac{g_{1,t}(Z_t \mid \mathcal{F}_{t-1})}{g_{0,t}(Z_t \mid \mathcal{F}_{t-1})}. \tag{33}$$

*Then $R_t$ is a conditional e-value increment under $M_0$, and $\prod_{u \leq t} R_u$ is an e-process.*

*Proof.* Conditional on $\mathcal{F}_{t-1}$, $Z_t$ has density $g_{0,t}(z \mid \mathcal{F}_{t-1})$ under $M_0$. Therefore,

$$\mathbb{E}_0(R_t \mid \mathcal{F}_{t-1}) = \int_{\{g_0 > 0\}} g_{1,t}(z \mid \mathcal{F}_{t-1})\nu_t(\mathrm{d}z) \leq 1.$$

The product result follows as before. □

*Remark* 6 (Discrete and mixed features)*.* The feature-level theorem holds for any dominating measure $\nu_t$. If $Z_t$ is discrete (such as a count of barrier crossings or state transitions within a block), $g_{0,t}$ and $g_{1,t}$ are probability mass functions, and $\nu_t$ is the counting measure. For discrete features, standard probability integral transforms (PIT) are not uniform, but one can use randomized PITs to obtain uniform continuous residuals, or work directly with the discrete probability mass functions in the density-ratio increment $R_t$.

##### S3.3.1 Rosenblatt residuals and step-angle dependence

Feature-level e-values are especially useful for circular-linear diagnostics. Suppose the null model gives a joint predictive distribution for $Y_t = (L_t, \Theta_t)$. A sequential Rosenblatt transform can be defined by

$$U_t^L = F_{0,t}^L(L_t \mid \mathcal{F}_{t-1}), \tag{34}$$

$$U_t^\Theta = F_{0,t}^{\Theta|L}(\Theta_t \mid L_t, \mathcal{F}_{t-1}), \tag{35}$$

where the second conditional distribution respects the circular support of $\Theta_t$. Crucially, $F_{0,t}^L$ and $F_{0,t}^{\Theta|L}$ are cumulative distribution functions derived from the marginal observable predictive mixture $p_0(y_t \mid \mathcal{F}_{t-1})$ (by marginalizing over the latent states using filtered probabilities), not from a decoded state or state-conditioned distribution. Under a continuous correctly specified null predictive distribution, $(U_t^L, U_t^\Theta)$ is conditionally uniform on $[0,1]^2$. A copula diagnostic can therefore use

$$R_t^{\mathrm{cop}} = c_\rho(U_t^L, U_t^\Theta), \tag{36}$$

where $c_\rho$ is a copula density on $[0,1]^2$, with $\rho$ fixed from training or chosen predictably. Theorem 12 gives conditional e-validity because $c_\rho$ integrates to one under the uniform null. This construction targets residual step-angle dependence after the fitted HMM has already accounted for latent behavioural states.

*Remark* 7 (Circular data)*.* Turning angles live on the circle, so residual diagnostics should respect periodicity. One may work with circular distribution functions relative to a chosen origin, with probability integral transforms for von Mises or mixture von Mises predictive distributions, or with features such as $\cos(\Theta_t - \mu_t)$ whose null law is derived from the circular predictive distribution. The e-value requirement is not normality of residuals; it is knowledge or independent estimation of the feature's null predictive law.

##### S3.3.2 Blockwise diagnostics for long-horizon movement summaries

One-step predictive diagnostics are local. Movement-model failures can also appear only over longer horizons, for example through utilization distributions, mean squared displacement, straightness, return times, residence times or barrier crossing frequencies. These can be handled by blockwise e-values.

**Theorem 13** (Blockwise predictive e-values for predictable boundaries)**.** *Let $0 = \tau_0 < \tau_1 < \tau_2 < \cdots$ be stopping times such that each block length $T_j = \tau_j - \tau_{j-1}$ is $\mathcal{F}_{\tau_{j-1}}$-measurable. This includes deterministic block boundaries (e.g., daily, weekly, or seasonal blocks). For block $j$, write $Y_{B_j} = Y_{\tau_{j-1}+1:\tau_j}$. Let $p_{0,j}(y_{B_j} \mid \mathcal{F}_{\tau_{j-1}}) = \prod_{t=\tau_{j-1}+1}^{\tau_j} p_0(y_t \mid \mathcal{F}_{t-1})$ be the conditional null predictive density of the block, and let $q_j(y_{B_j} \mid \mathcal{F}_{\tau_{j-1}})$ be a predictable diagnostic subdensity on $\mathcal{Y}^{T_j}$. Define*

$$E_j^{\text{block}} = \frac{q_j(Y_{B_j} \mid \mathcal{F}_{\tau_{j-1}})}{p_{0,j}(Y_{B_j} \mid \mathcal{F}_{\tau_{j-1}})}. \tag{37}$$

*Then $(\prod_{r \le j} E_r^{\text{block}})_{j \ge 0}$ is an e-process under $M_0$.*

*Proof.* Conditional on $\mathcal{F}_{\tau_{j-1}}$, the block $Y_{B_j}$ has density $p_{0,j}$ under $M_0$ and the length $T_j$ is fixed. Since $q_j$ is a subdensity on $\mathcal{Y}^{T_j}$, the same density-ratio calculation as in Theorem 1 of the main article gives $\mathbb{E}_0(E_j^{\text{block}} \mid \mathcal{F}_{\tau_{j-1}}) \le 1$. The cumulative product over blocks is therefore a nonnegative supermartingale with respect to the block filtration $(\mathcal{F}_{\tau_j})_{j \ge 0}$. □

**Theorem 14** (Blockwise e-values for genuinely stopped blocks)**.** *Suppose the block boundary $\tau_j$ is a stopping time that depends on observations inside the block (e.g., stopping when the animal leaves a home range or crosses a spatial boundary). Let the stopped block variable be $X_j = (Y_{B_j}, M_j)$, where $M_j = \tau_j - \tau_{j-1}$ is the random block length. Under $M_0$, the conditional joint density of the path $y_{1:m}$ and stopping event $M_j = m$ is:*

$$\begin{aligned} p_{0,j}(y_{1:m}, m \mid \mathcal{F}_{\tau_{j-1}}) = &\left( \prod_{t=\tau_{j-1}+1}^{\tau_{j-1}+m} p_0(y_t \mid \mathcal{F}_{t-1}) \right) \\ &\times \mathbb{P}_0(\tau_j - \tau_{j-1} = m \mid Y_{\tau_{j-1}+1:\tau_{j-1}+m} = y_{1:m}, \mathcal{F}_{\tau_{j-1}}). \end{aligned} \tag{38}$$

*Let $q_j(y_{1:m}, m \mid \mathcal{F}_{\tau_{j-1}})$ be a predictable diagnostic subdensity on the variable-length space $\bigcup_{m=1}^{\infty}(\mathcal{Y}^m \times \{m\})$. Define*

$$E_j^{\text{block}} = \frac{q_j(Y_{B_j}, M_j \mid \mathcal{F}_{\tau_{j-1}})}{p_{0,j}(Y_{B_j}, M_j \mid \mathcal{F}_{\tau_{j-1}})}. \tag{39}$$

*Then $(\prod_{r \le j} E_r^{\text{block}})_{j \ge 0}$ is an e-process under $M_0$.*

*Proof.* Conditional on $\mathcal{F}_{\tau_{j-1}}$, the random variable $X_j$ has joint density $p_{0,j}$ with respect to the product of the dominating measure on $\mathcal{Y}^m$ and the counting measure on $\mathbb{N}^+$. Since $q_j$ is a subdensity on the Stopped-Path space, we have:

$$\mathbb{E}_0(E_j^{\text{block}} \mid \mathcal{F}_{\tau_{j-1}}) = \sum_{m=1}^{\infty} \int \frac{q_j(y_{1:m}, m \mid \mathcal{F}_{\tau_{j-1}})}{p_{0,j}(y_{1:m}, m \mid \mathcal{F}_{\tau_{j-1}})} p_{0,j}(y_{1:m}, m \mid \mathcal{F}_{\tau_{j-1}}) \,\mathrm{d}\mu^{(m)}(y_{1:m}) \tag{40}$$

$$= \sum_{m=1}^{\infty} \int q_j(y_{1:m}, m \mid \mathcal{F}_{\tau_{j-1}}) \,\mathrm{d}\mu^{(m)}(y_{1:m}) \le 1. \tag{41}$$

The cumulative product over blocks is therefore a nonnegative supermartingale with respect to $(\mathcal{F}_{\tau_j})_{j\geq 0}$. □

**Corollary 15** (Feature-level block diagnostics). *Let $Z_j = h_j(Y_{B_j}, \mathcal{F}_{\tau_{j-1}})$ be a predictable block feature such as a barrier-crossing indicator, a displacement statistic or a summary of space use over the block. If the conditional null density or mass function of $Z_j$ is $g_{0,j}$ and a diagnostic alternative is $g_{1,j}$, then $g_{1,j}(Z_j)/g_{0,j}(Z_j)$ is a blockwise e-value increment.*

*Proof.* Apply Theorem 12 at the block level. □

*Remark* 8 (Simulation-based block laws). When the null law of a block feature is unavailable analytically, it may be approximated by simulation from the fitted HMM using only training information and the observed block covariates. Exact finite-sample e-validity then depends on how the density or mass estimate is constructed. A conservative recommendation for a first implementation is to fit any feature-level diagnostic law on independent null simulations or training data, and to reserve a separate validation trajectory for the final e-process calculation.

### S4 Simulation allocation and DGP details

#### S4.1 Simulation allocation

The main article reports the scenarios that directly support the central methodological narrative: calibration, targeted movement diagnostics, predictable localization and predictable diagnostic combination. This supplement records additional scenarios that are useful for sensitivity analysis, implementation warnings or methodological extensions.

#### S4.2 Simulation DGP details

To ensure the simulation results can be understood independently, we briefly outline the data-generating processes (DGPs) used in the main scenarios. Von Mises distributions in the simulations are parametrized through circular standard deviations (sd); the corresponding concentration parameters $\kappa$ were obtained by numerical inversion of the circular variance formula $V = 1 - I_1(\kappa)/I_0(\kappa)$, where $I_p$ denotes the modified Bessel function of order $p$. All parameters are specified on the natural scale.

- **S1 (Calibration)**: A 2-state HMM with initial probabilities $(0.5, 0.5)$ and transition matrix $\Gamma$ with $\Gamma(1,1) = 0.8$, $\Gamma(2,2) = 0.7$. State 1 (encamped) has step lengths drawn from Gamma$(\text{shape} = 2, \text{rate} = 0.5)$ and turning angles from von Mises$(\mu = 0, \text{sd} = 0.5)$. State 2 (exploratory) has step lengths Gamma$(10, 2)$ and turning angles von Mises$(\mu = 0, \text{sd} = 2)$. The diagnostic $q_t$ is an alternative 2-state HMM with slightly different parameters (e.g., transition and state-dependent parameters are perturbed relative to the null generator $p_0$).

| Scenario | Placement | Reason |
|---|---|---|
| S1 | Main article | Basic null calibration is required to support the e-process implementation. |
| S2 | Supplement | Parameter estimation uses an oracle-state estimator in the current simulation, so it is best treated as a sensitivity check. |
| S3 | Main article | The missing-state diagnostic is a core example of a full predictive HMM alternative. |
| S4 | Main article | Angular misspecification is a central movement-specific diagnostic. |
| S5 | Main article | Residual step-angle dependence is a central feature-level diagnostic. |
| S6 | Main article | Duration misspecification is a key movement failure and illustrates blockwise observable features. |
| S7 | Main article | Localization is a main theoretical and practical contribution. |
| S8 | Main article | Predictable mixtures and switching are main combination tools. |
| S9 | Supplement | The product example is an important warning, but it is not part of the recommended procedure. |
| S10 | Supplement | Long-horizon straightness is an advanced blockwise extension beyond the main simulation narrative. |
| S11 | Supplement | Individual cross-fitting is a validation-design extension. |
| S12 | Supplement | Composite envelopes are conservative and optional for the first article. |

Table S5: Allocation of simulation scenarios between the main article and supplementary material.

- **S3 (Missing state)**: The true DGP is a 3-state HMM with an intermediate speed state, but the null model is restricted to $K = 2$. The transition matrix is

$$\Gamma = \begin{pmatrix} 0.91 & 0.06 & 0.03 \\ 0.08 & 0.87 & 0.05 \\ 0.08 & 0.10 & 0.82 \end{pmatrix}.$$

  Step shapes are $(2.0, 7.5, 4.0)$, rates $(3.0, 2.0, 1.2)$, angle means $(0, 0, 0.6)$, and angle SDs $(1.6, 0.35, 0.75)$. The diagnostic $q_t$ uses the *true* 3-state DGP parameters *(oracle diagnostic)*.

- **S4 (Angular misspecification)**: The true DGP has step shapes $(2.0, 7.0)$, rates $(3.0, 2.0)$, and angle SDs $(1.8, 0.25)$. The null model is misspecified with angle SDs $(0.85, 0.30)$ and state 2 centered at $\mu = 0.25$ rather than 0. The diagnostic $q_t$ is a fixed, manually chosen diffuse-angle alternative *(fixed alternative)*.

- **S5 (Copula dependence)**: The true DGP includes a Gaussian copula with correlation $\rho = 0.6$ between step lengths and turning angles within each state, violating the conditional independence assumption of standard HMMs. The copula diagnostic uses a fixed target correlation $\rho_q = 0.55$ *(fixed alternative)*.

- **S6 (Duration)**: The true DGP is a hidden semi-Markov model (HSMM) where the dwell time in both states follows a shifted Poisson distribution (mean 18), rather than a geometric distribution. The duration diagnostic is a blockwise observable feature (switch counts within each block) and is *fixed* before seeing the data *(fixed alternative)*.
- **S7 (Localized failure)**: Similar to S5 ($\rho = 0.8$ in state 2) but restricted to a time window (times 121 to 220). The diagnostic $q_t$ uses a fixed predictable weight concentrated on the failure window *(fixed alternative)*.
- **S8 (Mixture switching)**: The alternative has two successive defects: angular misspecification (times 1 to 149) followed by step-angle copula dependence (times 150 to 300). The mixture and switching diagnostics combine fixed alternatives specified before the run *(fixed alternatives)*.

Future work could further characterize diagnostic sensitivity by exploring weaker alternatives, such as scenarios with highly overlapping state-dependent distributions, only slightly misspecified turning angles (e.g., small differences in concentration parameters), moderate copula dependence ($\rho \approx 0.1$), or near-geometric dwell time distributions.

### S5 Supplementary scenarios

#### S5.1 S2: train/validation with estimated parameters

Scenario S2 compares known parameters, oracle-state estimated parameters evaluated under the true generator and oracle-state estimated parameters evaluated under the fitted generator. At $\alpha = 0.05$, the crossing rates are 0.057, 0.053 and 0.023, respectively. These results are compatible with the split-validation interpretation, but the estimator uses simulated latent states. For that reason, S2 is not used as a main article result.

| Setting | Crossing rate | Monte Carlo SE | Mean final log e-value |
|---|---|---|---|
| Known parameters, true generator | 0.057 | 0.013 | -23.868 |
| Oracle-state estimate, true generator | 0.053 | 0.013 | -23.885 |
| Oracle-state estimate, fitted generator | 0.023 | 0.009 | -24.565 |

Table S6: S2 results at $\alpha = 0.05$.

#### S5.2 S9: warning against naive parallel products

Scenario S9 demonstrates the consequence of multiplying diagnostics computed in parallel on the same validation observations. At $\alpha = 0.05$, a single valid e-process and the weighted average of two copies both cross at rate 0.030, while naive products of two and three copies cross at rates 0.117 and 0.215. This supports the recommendation to use predictable mixtures, switching or weighted averages rather than parallel products.

| Method | Status | Crossing rate | Crossing rate minus $\alpha$ |
|---|---|---|---|
| Single e-process | Valid | 0.030 | -0.020 |
| Weighted average of two copies | Valid | 0.030 | -0.020 |
| Naive product of two copies | Not valid by default | 0.117 | 0.067 |
| Naive product of three copies | Not valid by default | 0.215 | 0.165 |

Table S7: S9 results at $\alpha = 0.05$.

#### S5.3 S10: blockwise long-horizon straightness

Scenario S10 uses block straightness, defined as net displacement divided by total path length, to detect a long-horizon failure generated by autocorrelated turning angles. At $\alpha = 0.05$, the crossing rate is 0.036 under the HMM null and 0.796 under the long-horizon alternative. This scenario supports the blockwise theory but is kept in the supplement because it is a more advanced extension.

| Generator | Crossing rate | Monte Carlo SE | Mean final log e-value | Mean straightness |
|---|---|---|---|---|
| HMM null | 0.036 | 0.012 | -4.909 | 0.496 |
| Autocorr. angle alternative | 0.796 | 0.025 | 4.639 | 0.347 |

Table S8: S10 results at $\alpha = 0.05$.

#### S5.4 S11: individual validation and cross-fitted averages

Scenario S11 studies validation by individuals. The weighted average of final individual e-values is conservative under the true and fitted null settings, with crossing rates 0.000 and 0.004 at $\alpha = 0.05$. The exploratory scan over any individual is not a globally valid e-value and crosses at rates 0.304 and 0.228 under the two null settings. Under a single failed individual, the cross-fitted average crosses at rate 0.996 and the individual scan crosses at rate 1.000.

| Scenario | Method | Status | Crossing rate |
|---|---|---|---|
| True null | Cross-fitted final average | Valid final e-value | 0.000 |
| True null | Any individual scan | Exploratory | 0.304 |
| Fitted null | Cross-fitted final average | Valid final e-value | 0.004 |
| Fitted null | Any individual scan | Exploratory | 0.228 |
| Single failed individual | Cross-fitted final average | Valid final e-value | 0.996 |
| Single failed individual | Any individual scan | Exploratory | 1.000 |

Table S9: S11 results at $\alpha = 0.05$.

#### S5.5 S12: finite-family composite envelope

Scenario S12 studies a conservative envelope over a finite family of three HMM null generators. It is important to note that the practical use of the composite envelope (Proposition 2) is primarily intended for finite or compactly restricted families. Continuous envelopes of unbounded parameter families for densities such as Gamma or von Mises can become numerically unstable or computationally prohibitive. The center-only denominator is not uniform over the family and gives a crossing rate of 0.980 when the true generator is the diffuse-angle null. The composite envelope gives crossing rate 0.000 at $\alpha = 0.05$ for each generator in the finite family. The scenario illustrates uniform validity and its cost in conservatism.

| Generator | Method | Status | Crossing rate |
|---|---|---|---|
| Center null | Oracle true null | Valid for known generator | 0.020 |
| Center null | Center-only denominator | Not uniform over family | 0.020 |
| Center null | Composite envelope | Uniform finite-family envelope | 0.000 |
| Persistent null | Oracle true null | Valid for known generator | 0.027 |
| Persistent null | Center-only denominator | Not uniform over family | 0.000 |
| Persistent null | Composite envelope | Uniform finite-family envelope | 0.000 |
| Diffuse-angle null | Oracle true null | Valid for known generator | 0.043 |
| Diffuse-angle null | Center-only denominator | Not uniform over family | 0.980 |
| Diffuse-angle null | Composite envelope | Uniform finite-family envelope | 0.000 |

Table S10: S12 results at $\alpha = 0.05$.

### S6 Comparison with classical pseudo-residual diagnostics

The main article compares the proposed predictive e-diagnostics against classical diagnostic tests based on pseudo-residuals. Componentwise marginal probability integral transform (PIT) residuals are distinguished from joint sequential Rosenblatt residuals: under a correctly specified movement HMM, the sequential PIT residuals computed from the full one-step conditional predictive distribution for step lengths and turning angles individually form sequences of independent Uniform$(0, 1)$ variables, but these componentwise marginal PITs do not capture contemporaneous dependence between the two dimensions within the same step. To obtain fully independent Uniform$(0, 1)$ variables across both dimensions and time, one constructs joint sequential Rosenblatt residuals $U_t^L = F_{0,t}^L(L_t \mid \mathcal{F}_{t-1})$ and $U_t^\Theta = F_{0,t}^{\Theta|L}(\Theta_t \mid L_t, \mathcal{F}_{t-1})$ and maps them to standard normal variables to perform standard tests (Zucchini et al., 2016). The classical tests are retrospective terminal tests evaluated at the end of the validation trajectory and are not adjusted for sequential or multiple checking, so their power is not directly comparable but serves as a useful retrospective benchmark. The three classical tests considered are a Kolmogorov-Smirnov (KS) test on the circular PIT turning-angle residuals and the marginal step-length PIT residuals, a Ljung-Box test on the normal step pseudo-residuals at lag 5, and a correlation test on the normal step and angle pseudo-residuals.

Figure S3 summarizes the empirical signal and rejection rates across the simulated trajectories, using the scenario-specific validation length reported in the main article ($T = 300$ for S3–S6).

1. **Underfitted states (S3)**: the e-diagnostic detects the missing third state in 99.7% of the replicates, whereas the classical marginal KS tests on steps and angles fail to reject

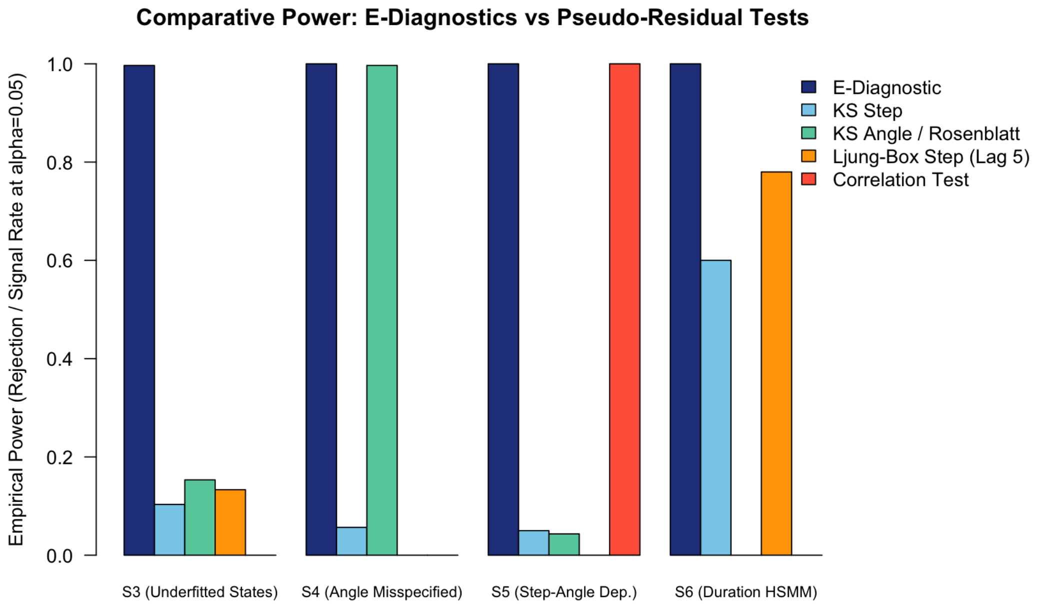


Figure S3: Signal/rejection rates in scenarios S3–S6. The validation length is $T = 300$ in all four scenarios; S3–S5 use 300 replicates and S6 uses 250 replicates.

the null in most replicates (10.3% and 15.3% power, respectively), and the Ljung-Box autocorrelation test achieves only 13.3% power.

2. **Angular misspecification (S4)**: both the angular e-diagnostic and the circular-linear KS test on turning angles exhibit near-perfect power (100% vs. 99.7%).

3. **Step-angle joint dependence (S5)**: the copula e-diagnostic detects the residual dependence in 100% of the replicates. The marginal KS tests fail (5.0% and 4.3% power) because they target univariate deviations only. The targeted correlation test on the joint normal residuals also achieves 100% power, but the e-diagnostic achieves this without sacrificing sequential validity.

4. **Non-geometric state durations (S6)**: the blockwise duration e-diagnostic achieves 100% power in detecting the HSMM-like state transitions, whereas the step-length KS test and Ljung-Box test reach only 60.0% and 78.0% power, respectively.

These results show targeted sensitivity to the structured misspecifications used in these controlled scenarios. The comparison does not imply a universal dominance of e-diagnostics over classical tests, especially since the classical tests are evaluated retrospectively without adjustment for sequential monitoring or multiple testing.

| Diagnostic target | Null component | Diagnostic alternative | Ecological interpretation |
|---|---|---|---|
| Number of states | $K$-state HMM | $K+1$ state HMM or richer latent structure | Detects behaviour classes merged by the null model |
| Angular distribution | State-specific von Mises or wrapped distribution | Mixture of von Mises, flexible circular density or circular residual tilt | Detects multimodality, asymmetry or wrong concentration of turning angles |
| Step-angle dependence | Conditional independence $f^L_{0,s}(l)f^\Theta_{0,s}(\theta)$ | Joint circular-linear density or copula on predictive residuals | Detects whether speed and directionality remain dependent within states |
| State durations | Geometric dwell times implied by HMM | HSMM or duration-aware predictive model | Detects behaviours that persist too long or too briefly for a Markov state process |
| Temporal nonstationarity | Time-homogeneous parameters | Season-, phase- or covariate-varying diagnostic alternative | Detects periods where the fitted generator stops predicting well |
| Individual heterogeneity | Shared parameters or random effects in null | Individual-specific or hierarchical diagnostic alternative | Detects held-out animals poorly represented by the fitted population model |
| Long-horizon spatial behaviour | One-step HMM generator | Blockwise features such as displacement, barrier crossing, residence or space-use summaries | Detects failures visible only through simulated trajectories or aggregated movement behaviour |

Table S11: Movement-specific predictive e-diagnostics.

### S7 Movement-specific diagnostic catalogue

Table S11 summarizes a practical menu of diagnostics. Each diagnostic can be implemented as a full predictive density ratio, a feature-level e-value, or a blockwise e-value depending on the available null law and the scientific target.

#### S7.1 Diagnostic for the number of states

A natural alternative to a $K$-state HMM is a $(K+1)$-state HMM. Let $p_K(y_t \mid \mathcal{F}_{t-1})$ denote the observable predictive density of the $K$-state null HMM (i.e., $p_0$) and let $p_{K+1}(y_t \mid \mathcal{F}_{t-1})$ denote the predictive density of a $(K+1)$-state alternative. The ratio

$$E_t^{K+1} = \frac{p_{K+1}(Y_t \mid \mathcal{F}_{t-1})}{p_K(Y_t \mid \mathcal{F}_{t-1})}, \tag{42}$$

where both predictive densities marginalize over their respective latent states. If $\log E_{1:t}^{K+1}$ grows only during certain seasons or individuals, the diagnostic suggests that an additional behavioural mode is needed only locally rather than globally.

#### S7.2 Diagnostic for angular misspecification

Many movement HMMs use a state-specific von Mises distribution for turning angles. A diagnostic alternative can replace each state-specific angular density by a mixture of von Mises distributions, a wrapped distribution with heavier tails, or a feature-level e-value based on circular residuals. The purpose is not necessarily to adopt the alternative as the final model, but to test whether the null predictive distribution of turning angles can survive a targeted circular bet.

#### S7.3 Diagnostic for residual step-angle dependence

A common HMM assumption is conditional independence of step length and turning angle within state:

$$f_{0,s}(l, \theta) = f^{L}_{0,s}(l) f^{\Theta}_{0,s}(\theta). \tag{43}$$

A copula-based alternative uses

$$f_{1,s}(l, \theta) = c_s\{F^{L}_{0,s}(l), F^{\Theta}_{0,s}(\theta); \rho_s\} f^{L}_{0,s}(l) f^{\Theta}_{0,s}(\theta), \tag{44}$$

with parameters estimated on training data or selected predictably. A feature-level version applies the Rosenblatt residual construction described in Section S3.3. This diagnostic asks whether, after accounting for latent behaviour, longer steps are associated with different directional persistence or turning behaviour.

#### S7.4 Diagnostic for duration misspecification

A standard HMM implies geometric dwell times in latent states. HSMMs relax this by allowing arbitrary state dwell-time distributions (Langrock and Zucchini, 2011). A duration diagnostic may use an HSMM predictive density as $q_t$, or a blockwise feature that summarizes likely persistence in behaviours. Because states are latent, duration diagnostics should be based on the predictive density of observations under the duration-aware alternative, not on treating decoded durations as observed without uncertainty.

#### S7.5 Diagnostic for long-term generative behaviour

Some movement failures are not local in $Y_t = (L_t, \Theta_t)$. A model can predict one-step movement well but fail to reproduce utilization distributions, corridors, barriers or returns. The blockwise theory in Theorems 13 and 14 supports diagnostics based on multi-step predictive distributions or block features. For example, for a monthly block one may define $Z_j$ as a barrier-crossing count or displacement statistic. A diagnostic alternative $g_{1,j}$ can then be compared with the null block law $g_{0,j}$ through $g_{1,j}(Z_j)/g_{0,j}(Z_j)$.

### S8 Recommended implementation workflow

A first implementation should prioritize exact split-sample validity and transparent diagnostics.

**Algorithm: predictive e-diagnostics for a movement HMM.**

1. Split the data into training and validation components. For animal movement, a held-out-individual split is often preferable when several individuals are available.

2. Fit the null HMM $M_0$ on the training data. Record all preprocessing, covariate choices and decisions regarding the number of states in $\mathcal{H}$.

3. Define a diagnostic menu before validation. Examples include $K + 1$ states, angular flexibility, step-angle dependence, duration structure and blockwise spatial summaries.

4. For each validation time $t$, compute the null observable predictive density by filtering:

$$p_0(Y_t \mid \mathcal{F}_{t-1}) = \sum_{s=1}^{K} a_{t|t-1}(s) f_{0,t,s}(Y_t \mid \mathcal{F}^{\mathrm{cov}}_{t-1}).$$

5. Compute the diagnostic predictive density or feature density $q_t(Y_t \mid \mathcal{F}_{t-1})$ for each chosen diagnostic.

6. Accumulate

$$\log E_{1:t} = \sum_{u=1}^{t} \{\log q_u(Y_u \mid \mathcal{F}_{u-1}) - \log p_0(Y_u \mid \mathcal{F}_{u-1})\}.$$

7. Monitor the anytime-valid threshold $\log(1/\alpha)$. For $\alpha = 0.05$, this threshold is $\log(20) \approx 2.996$.

8. Report both global and localized evidence: individual e-processes, soft state-localized processes, segment-specific processes and local increments.

9. Interpret the result comparatively. A large e-process indicates that the diagnostic alternative predicted the validation sequence better than the fitted HMM in a sequentially valid sense; it does not by itself establish an ecological mechanism.

### S9 Reproducibility

The simulation scripts are stored in the project directory `simulations/`. The complete simulation suite is run by `simulations/run_all_simulations.R`. The elk application is run by `application/run_application.R`. Generated outputs are written to:

- `results/simulation_tables/`;
- `results/simulation_figures/`;
- `results/application_tables/`;
- `results/application_figures/`;
- `paper/figures/`.

The public project repository at `https://github.com/AurelienNicosiaULaval/evalue-HMM` provides the code and documentation needed to reproduce the analyses. Generated outputs can be regenerated from the scripts; random seeds are set within the simulation scripts, and the elk application is deterministic conditional on the installed package versions and deterministic starting profiles.
